# Structural spine plasticity: learning and forgetting of odor-specific subnetworks in the olfactory bulb

**DOI:** 10.1101/2022.06.29.498211

**Authors:** John Hongyu Meng, Hermann Riecke

## Abstract

Learning to discriminate between different sensory stimuli is essential for survival. In rodents, the olfactory bulb, which contributes to odor discrimination via pattern separation, exhibits extensive structural synaptic plasticity involving the formation and removal of synaptic spines, even in adult animals. The network connectivity resulting from this plasticity is still poorly understood. To gain insight into this connectivity we present here a computational model for the structural plasticity of the reciprocal synapses between the dominant population of excitatory principal neurons and inhibitory interneurons. It incorporates the observed modulation of spine stability by odor exposure. The model captures the striking experimental observation that the exposure to odors does not always enhance their discriminability: while training with similar odors enhanced their discriminability, training with dissimilar odors actually *reduced* the discriminability of the training stimuli (Chu et al, 2016). Strikingly, this differential learning does not require the activity-dependence of the spine stability and occurs also in a model with purely random spine dynamics in which the spine density is changed homogeneously, e.g., due to a global signal. However, the experimentally observed odor-specific reduction in the response of principal cells as a result of extended odor exposure (Kato et al. 2012) and the concurrent disinhibition of a subset of principal cells arise only in the activity-dependent model. Moreover, this model predicts the experimentally testable recovery of odor response through weak but not through strong odor re-exposure and the forgetting of odors via exposure to interfering odors. Combined with the experimental observations, the computational model provides strong support for the prediction that odor exposure leads to the formation of odor-specific subnetworks in the olfactory bulb.

**Author Summary:** A key feature of the brain is its ability to learn through the plasticity of its network. The olfactory bulb in the olfactory system is a remarkable brain area whose anatomical structure evolves substantially still in adult animals by establishing new synaptic connections and removing existing ones. We present a computational model for this process and employ it to interpret recent experimental results. By comparing the results of our model with those of a random control model we identify various experimental observations that lend strong support to the notion that the network of the olfactory bulb comprises learned, odor-specific subnetworks. Moreover, our model explains the recent observation that the learning of odors does not always improve their discriminability and provides testable predictions for the recovery of odor response after repeated odor exposure and for when the learning of new odors interferes with retaining the memory of familiar odors.

## Introduction

Learning and long-term memory are to a large extent implemented through the plasticity of neuronal connectivity. In adult animals this plasticity is typically dominated by long-term potentiation and long-term depression, which change the strength of the synapses, while during development structural plasticity in the form of the addition and removal of neurons as well as the formation and pruning of synapses plays a central role. Strikingly, in olfaction structural plasticity appears to remain a key component of learning even in adult animals. Thus, new adult-born granule-cell interneurons are persistently integrated into the olfactory bulb, which is the first brain region to receive sensory input from the nose, and new synapses between granule cells and the principal mitral cells are formed. In parallel, granule cells undergo controlled apoptosis [1, 2], the extent of which, however, has recently come under dispute ([3], see also discussion with the reviewers in [4]), and synapses are removed [5, 6]. This makes the olfactory bulb an excellent system to study structural plasticity.

A widely assumed function of the olfactory bulb is to aid in the discrimination between odors by enhancing differences in the activity patterns that they evoke [7, 8]. Computational modeling has identified a number of mechanism by which this can be achieved, which are not mutually exclusive. This includes a nonlinear response of the neurons without any lateral interactions among the neurons [9] or with random [10] or all-to-all connectivity [11]. Given the extensive plasticity of the olfactory bulb, adaptive connectivity with linear or nonlinear response of the neurons has also been identified as an efficient mechanism for pattern separation [6, 12–15]. A common approach to probe this function of the olfactory system is to test whether animals learn to discriminate between two similar odors. In these tasks their correct choice may or may not be associated with a reward. It has been found that certain types of perceptual learning depend on the olfactory bulb [16], require adult neurogenesis [17], and are associated with increased densities of the spines on granule-cell (GC) dendrites that form the synapses with the mitral cells (MCs) [18]. The mechanisms controlling these plasticity mechanisms are not very well understood. The survival of GCs is enhanced by odor enrichment [4, 19] and by GC activity [20]. Similarly, odor enrichment increases the stability of spines in bulbar regions that are activated by the enrichment odors [5, 21]. Simultaneous measurements of the spine stability and the activity of the cells associated with that synapse are challenging. Thus, knowledge about the response of the bulbar spine dynamics to signals (like glutamate, BDNF) associated with neuronal activity is somewhat limited [22, 23]. So far no direct connection between the activity of individual neurons, the stability of their spines, and the circuits emerging from that plasticity has been established experimentally. In fact, the organizational principle of the network of mitral and granule cells is still only poorly understood, reflecting also the lack of significant chemotopical organization of the bulb [24]. The overall dependence of the spine stability on odor enrichment suggests that the resulting connectivity would reflect the specific odors that the animal has experienced. Computational modeling predicts such a structure and suggests that it could underly the observed improvement in odor discrimination [1, 2, 6].

In this paper we pursue two goals. Motivated by a number of experimental observations [6, 18, 22, 23], we first put forward a phenomenological model for the activity dependence of the spine dynamics that incorporates the memory of previous odor exposures. We then use this model to interpret two sets of experimental results [25, 26], focusing particularly on the question to what extent these experiments support the computational prediction that the olfactory bulb exhibits a learned, odor-specific subnetwork structure. In [25] the repeated exposure to an odor over multiple days has been found to lead to a reduction of the mitral cell response that is specific to that odor, i.e. the response of the mitral cell to a different odor changes only little during the same time. A different set of experiments [26] revealed that repeated odor exposure makes the bulbar mitral cell representations of two odors more discriminable if the two odors are very similar. However, if the two odors are very different from each other, the learning makes their representations actually less discriminable. Moreover, while the repeated odor exposure leads to an overall reduction of the mitral cell responses, individual mitral cells can exhibit an increase in activity and this increase can be odor-specific, i.e. for one of two very similar odors the mitral cell response can increase, while it remains low for the other odor, resembling differential disinhibition. Differential inhibition is also observed.

We show that our model is able to capture these observations. To disentangle the role of the activity-dependence of the spine dynamics we also consider a control model with purely random spine dynamics in which the statistically homogeneous spine density is modulated by a global signal. Somewhat surprisingly, the control model can capture the learning-induced improved discrimination of similar stimuli as well as the learning-induced reduced discrimination of dissimilar stimuli. However, the experimentally observed disinhibition of mitral cells is extremely unlikely in such a model. In combination with [25, 26] our model gives therefore strong support to the notion that the structural plasticity in the olfactory bulb leads to a learned, odor-specific network structure in which MCs preferentially inhibit each other disynaptically, if they have similar receptive fields [6, 12, 14, 15]. Moreover, the model makes testable predictions for the recovery of MC responses after repeated odor exposures, for the suppression of odor discrimination by odor exposure, and for the forgetting of odor memories through interference. An earlier account of some of these results is available at [27].

## Results

### Formulation of the model

Using computational modeling we investigated the ramifications of the structural plasticity of the reciprocal synapses between mitral cells (MCs) and granule cells (GCs). We focused on those two cell types and did not include the processing in the glomerular layer between the sensory neurons and the mitral cells. Instead, we considered the glomerular activation patterns as inputs to the MCs (Fig 1A). The MCs and GCs were described by firing-rate models. Reflecting the reciprocal nature of the MC-GC synapses the GCs inhibited those MCs that excited them as indicated in the inset of Fig 1A, which depicts a spine on a GC dendrite and an excitatory synapse (MC→GC) and an inhibitory synapse (GC→MC). Since the synapses are located on the secondary dendrites of the MCs, which reach across large portions of the olfactory bulb [28], we allowed synapses to be formed between all MCs and all GCs without any spatial limitations. We recognize that this ignores the sparse structure of the dendrites, which results in a decrease in the connection probability between widely separated MCs [29, 30].

**Fig 1.**
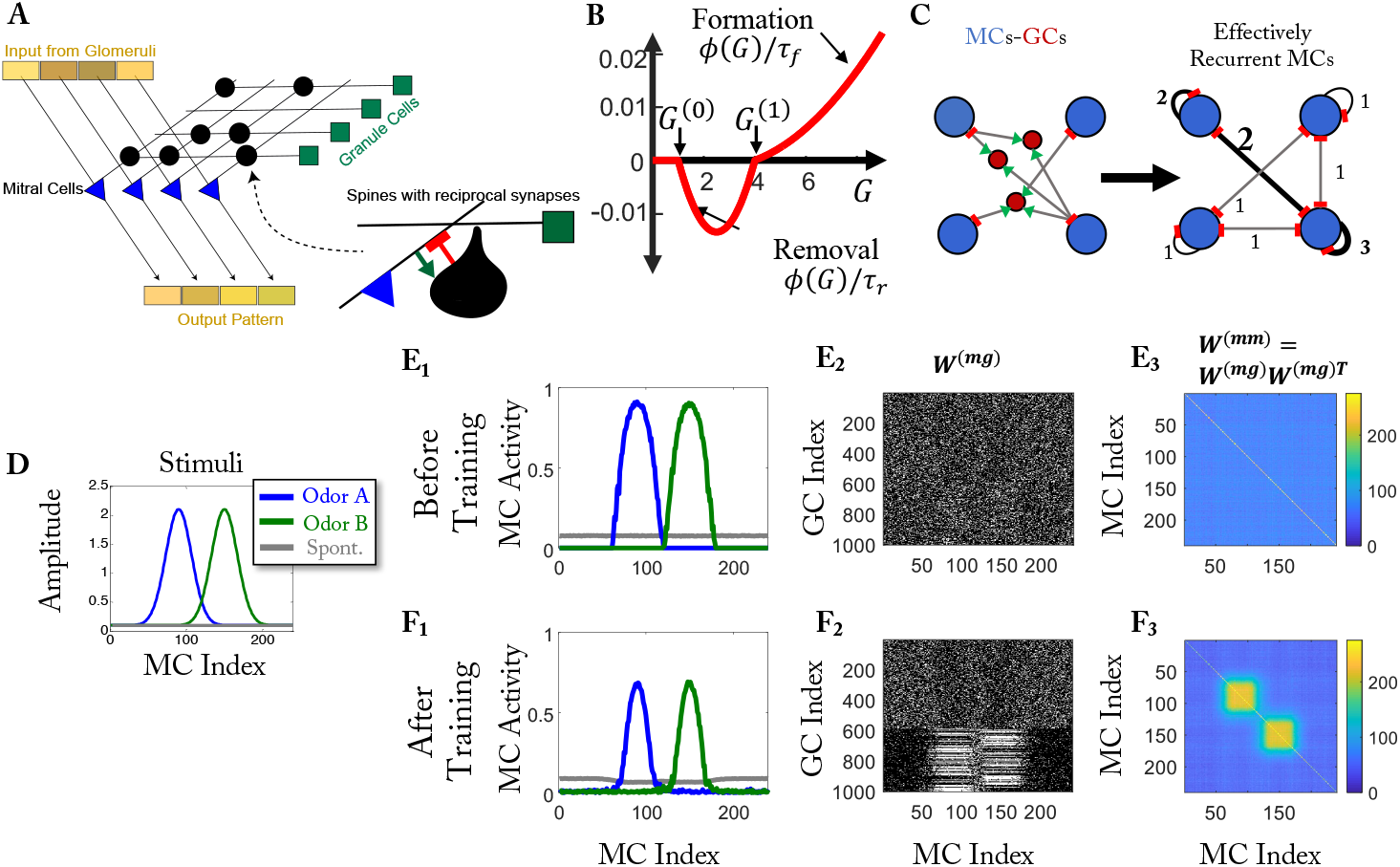
Computational model. (A) Sketch of the network. All spines provide reciprocal synapses that excite GCs (green arrow) and inhibit MCs (red bar). (B) Spine formation is controlled by *R* = *Mϕ*(*G*). The figure shows *ϕ/τ*_*f*_ for *ϕ >* 0 and *ϕ/τ*_*r*_ for *ϕ <* 0 (cf. Eq. 5). (C) Disynaptic recurrent inhibition of MCs via GCs. The numbers in the right panel indicate the strength of the effective inhibition. (D) Simplified training stimuli. (E_1_, F_1_) MC activity before and after training, respectively. (E_2_, F_2_) Connectivity between MCs and GCs before and after training, respectively. Each white dot represents a connection between an MC and a GC. (E_3_, F_3_) Effective recurrent connectivity among MCs. The color represents the number of GCs that mediate the mutual disynaptic inhibition of MCs via GCs (cf. C).

The MC-GC synapses do exhibit some synaptic-weight plasticity. Its nature is, however, still quite unclear [31, 32]. Moreover, during perceptual learning the frequency of the inhibitory postsynaptic currents received by the MCs from the GCs has been found to increase, while their amplitude remained unchanged [18], suggesting that only the number but not the strength of the synapses changed. We therefore assume all connections to have the same weight and focus on the structural plasticity of the synapses. Thus, in each time step we add and remove reciprocal synapses between MCs and GCs with probabilities that depend on the activity of the cells that are connected by that synapse.

Our modeling is qualitatively motivated by a number of experimental studies. The overall stability of spines has been observed to be enhanced if animals experienced an enriched odor environment [5]. More precisely, across multiple days, spines in bulbar areas that were activated by the enrichment odors were stabilized, while those in other areas were not. Very recently, a positive correlation between the stability of individual spines and the GC-activity of the dendrite on which they are located has been observed [33]. Since filopodia often are precursors of spines, we also glean information from experiments addressing their dynamics. The frequency of formation and removal of filopodia has been found to increase with NMDA-receptor activation [22] and to decrease with Mg^2+^-concentration. This suggests that the formation rate of spines increases with the activity of the GC-dendrite on which they are forming. The dynamics of filopodia that are located on spine heads are often indicative of subsequent dynamics of that spine. Specifically, the amplitude and direction of the movement of the spine head has been found to be correlated with the lifetime and orientation of such a filopodium, respectively [23]. Moreover, the dynamics of the filopodia have been found to be triggered by presynaptic glutamate release, suggesting a role of MC-activity in the spine dynamics [23]. Biophysical details about the mechanisms relating the spine dynamics with the temporal evolution of the neuronal activity during odor exposure are, however, not well known yet.

Based on these observations, we assume the formation and removal of synapses to follow Poisson processes with rates that depend on the activities of the neurons connected by the respective synapse (for details see Methods). We express the formation and removal rates in terms of a single rate function *R* = *Mϕ*(*G*) that depends on the activities *M* and *G* of the MC and the GC, respectively, that are connected by that synapse (Fig 1B). Similar to models for synaptic weight plasticity [34, 35], we take this function to be non-monotonic. For GC activities above a threshold *G*^(1)^ a new synapse is formed, while it is removed below that threshold as long as it is above a second, lower threshold, *G*^(0)^. For GC activities below *G*^(0)^ structural change is negligible. This lower threshold allows the synaptic connections of a GC to be remembered during times when the GC is not activated by any odor. Connections are removed, however, on GCs that are excited, but only weakly so. Since the change in the total number of synapses appears to be limited in experiments [18, 36], we hypothesize that there exists a homeostatic mechanism that keeps the total number of synapses on a given GC within a limited range. For most of our results, we employ a top-*k* competition mechanism, in which only the *k* strongest synapses survive. We show that qualitatively the same results are obtained with a more biophysical mechanism that is based on a limited resource (see Realization of competition through competition for a limited resource and Methods). Both mechanisms operate on a relatively fast time scale, as is generally needed for the stability of networks with Hebbian plasticity [37].

We are particularly interested in the impact of the plasticity on the ability of the network to learn to discriminate between stimuli that the network is repeatedly exposed to. We train the network by using a sequence of alternating stimuli A and B as sensory inputs. We assess the ability of the network to discriminate between these stimuli by comparing the difference in the MC activity patterns in response to the stimuli with the variability of these patterns across repeated presentations of the same stimuli. The firing rates themselves do not fluctuate between successive presentations. However, the spike trains underlying those mean firing rates can fluctuate. Assuming Poisson-like spike trains, we take the mean firing rate of an MC as a proxy for its variance (for details see Methods section). To illustrate the mechanism by which the discriminability is modified, we use simplified stimuli (Fig 1D), while naturalistic stimuli are used to compare with experimental data (Fig 2A).

**Fig 2.**
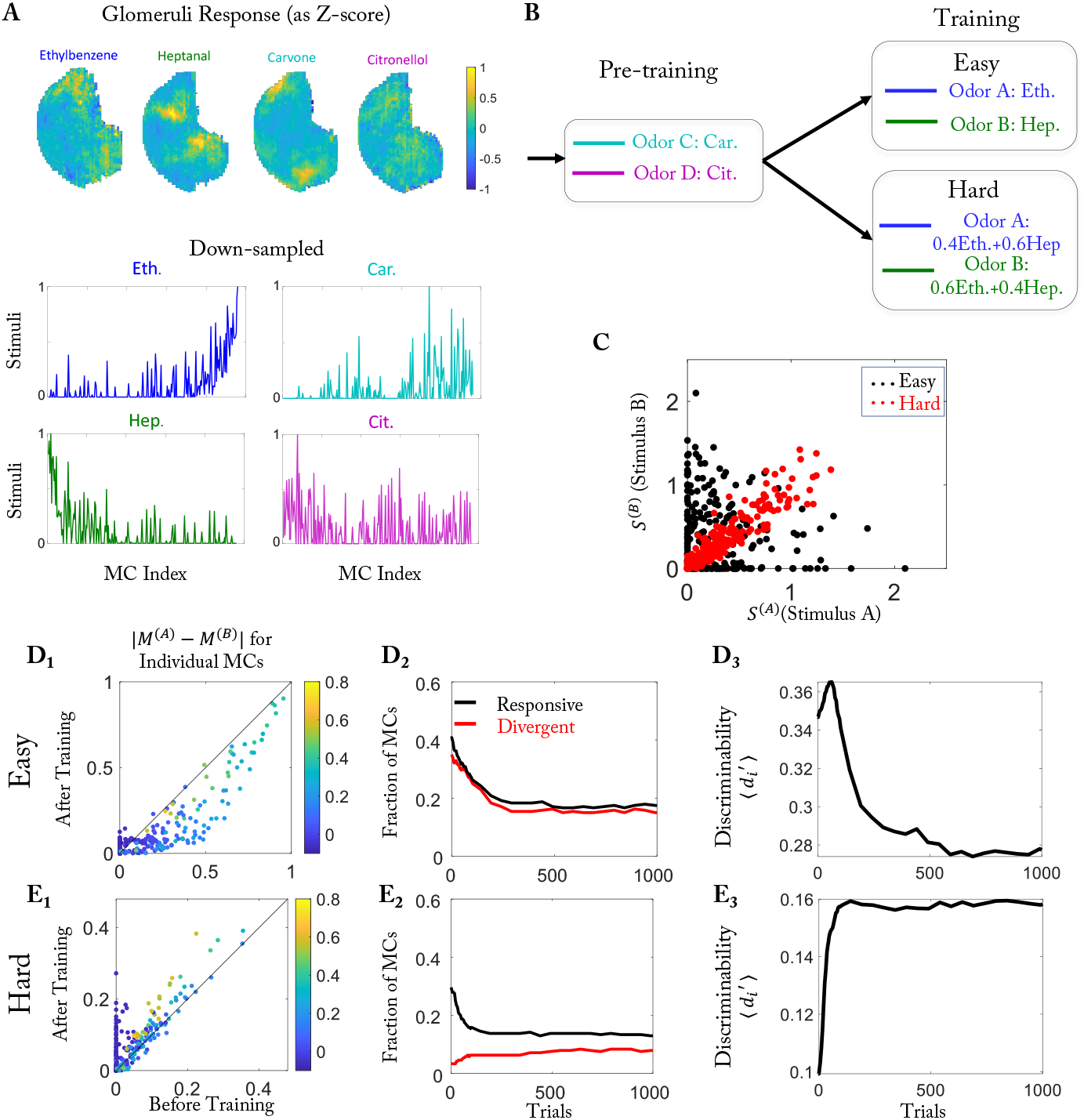
Easy and hard discrimination task using naturalistic stimuli. (A) Odors employed in the training. Top: Glomerular activation patterns (cf. [39]). Bottom: Activation patterns down-sampled to 240 points, serving as stimuli **S**^(*i*)^. MCs are sorted based on the difference in activation by the stimuli of odor pair 1. (B) Computational protocol. (C) Activity of each MC for the two stimuli used in the easy (black) and hard (red) task. (D_1_,E_1_) Difference in the response to the two stimuli for each MC before and after training for the easy and the hard task, respectively. Color of the dots represents the mean response |***M*** ^(1)^ + ***M*** ^(2)^ - 2***M*** ^(*air*)^|*/*2 before training. (D_2_,E_2_) Temporal evolution of the number of responsive MCs and divergent MCs during the training in the easy task and hard task, respectively (cf. [26]). (D_3_,E_3_) Temporal evolution of 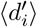 for the easy task and the hard task, respectively. Parameters as in Table 1 except for *γ* = 1.7 · 10^*-*4^.

### Plasticity induces mutual disynaptic inhibition of co-activated MCs

To illustrate the network evolution resulting in the structural-plasticity model, we use a pair of simple model stimuli A and B that excite partially overlapping sets of MCs (Fig 1D). In the absence of any odors (‘air’) the input is taken to be non-zero and homogeneous, reflecting the spontaneous activity of the MCs or of the neurons driving their input. Initially, the MC-GC connectivity is chosen to be random (Fig 1E_2_). The learning process reflects the effectively Hebbian character of the structural plasticity rule through which connections are established between cells that fire together. If a GC responds to a stimulus by integrating excitatory inputs from activated MCs, other MCs that are activated but not yet connected to this GC are likely to form a connection with that GC. This is reflected by the two blocks in Fig 1F_2_ involving GCs with indices above ∼ 600. Each of these GCs is connected with either the MCs responding to stimulus A (MC indices 60-110) or to stimulus B (MC indices 110-160). Conversely, if two MCs respond to the same stimulus, the number of GCs they both connect to increases, if the model is trained by that stimulus. Due to the reciprocal nature of these synapses, this induces enhanced mutual disynaptic inhibition (Fig 1C) between these MCs. This is seen in the two blocks along the diagonal line of their effective connectivity matrix, which gives the number of GCs that mediate that inhibition (Fig 1F_3_). As a result, the training preferentially reduces the activity of initially highly activated MCs (Fig 1E_1_, F_1_), consistent with observations [25, 38].

**Table 1.** Parameters used in the simulations of the model (unless stated otherwise).

|  |  |  |  |
| --- | --- | --- | --- |
| $N_{MC}$ | 240 | $N_{GC}$ | 1000 |
| $N_{conn}$ | 60 | $k$ | 66 |
| $\tau_M$ | 1 | $\tau_G$ | 0 |
| $\Delta t$ | 1 | | |
| $g_{thr}$ | 4.4 | $\gamma$ | $5e - 4$ |
| $G^{(0)}$ | 1 | $G^{(1)}$ | 4 |
| $\lambda_f$ | $6 \cdot 10^{-4}$ | $\lambda_r$ | $6 \cdot 10^{-3}$ |

### Training enhances or reduces stimulus discriminability depending on their similarity

Intriguingly, in mice learning to discriminate between two odors the training enhanced the discriminability of the training stimuli *only* if they were similar, i.e., for hard discrimination tasks [26]. If the task was easy, i.e. if the stimuli were dissimilar, the training actually *reduced* the discriminability of the training stimuli. Here discriminability was measured using d’, which is given by the mean of the difference in MC-activity in response to the two odors relative to the square-root of the pooled variance. To provide insight into this surprising finding we employed the same training protocol as in the experiment, using naturalistic stimuli that were adapted from glomerular activation data of the Leon lab [39] (Fig 2A (top)) and down-sampled to reduce the computational effort (Fig 2A (bottom)). These stimuli were used for the easy discrimination task. As in [26], we used mixtures of the same stimuli for the hard discrimination task (60% : 40% *vs*. 40% : 60%, see Methods). The two odors of the hard task drove each MC to a very similar degree (Fig 2C).

Our model successfully reproduced several experimental observations of [26]. There it was found that training with the easy task *reduced* the difference in the response of the MCs to the two odors used in the task; the majority of the MCs fell below the diagonal in Fig 2D_1_. Here the response was defined as the difference between odor-evoked activity and air-evoked activity and the color of the dots in Fig 2D_1_,E_1_ denotes the average response of the respective MC across the two odors before training. In contrast, the difference *increased* with training in the hard task (Fig 2 E_1_). In particular, a fraction of cells showed the same response to both odors before training, but significantly different responses after training (Fig 2E_1_, dots along the y-axis). As in [26], we quantified these changes by classifying MCs into responsive cells, i.e., cells that showed a significant response to at least one of the two odors, and divergent cells, for which the two odors evoked significantly different responses (see Methods). In the easy task, not only the number of responsive cells decreased but also the number of divergent cells (Fig 2D_2_). In the hard task the number of responsive cells also decreased. The number of divergent cells, however, actually increased by a small amount (Fig 2E_2_). Thus, both results recapitulated the experimental observations [26]. These outcomes were not sensitive to the threshold *θ* that classified the cells (Fig S14).

The discriminability of two neuronal activity patterns depends not only on the difference of their mean firing rates but also on the trial-to-trial variability of their spike trains. The latter is not included in our firing-rate framework. It is quite common for the variability to increase with the firing rate; for instance, for Poisson spike trains the variance in the spike count is equal to the mean spike count, i.e. the variability increases like the square-root of the mean. As a proxy for the discriminability of the response patterns, we therefore defined 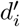 of each divergent neuron as the difference of the mean activity in response to the two odors divided by the square-root of the sum of the means (see Eq.(12) in the Methods). As in [26], training increased the mean 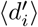 across the MCs in the hard task (Fig 2E_3_), but it decreased 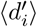 in the easy task (Fig 2D_3_), indicating reduced discriminability.

### Visualization of reduction and enhancement of discriminability

To better understand and visualize the mechanism underlying the change in discriminability, we employed simplified stimuli (Fig 3A,B). As in the case of realistic stimuli, the stimuli for the hard task were mixtures of two easily discriminated stimuli. Again, we first exposed the network to a set of pre-training stimuli. They set up an initial connectivity that was independent of the task stimuli (Fig 3A). It affected the initial response of the network to the training stimuli (Fig 3C_1_,D_1_).

**Fig 3.**
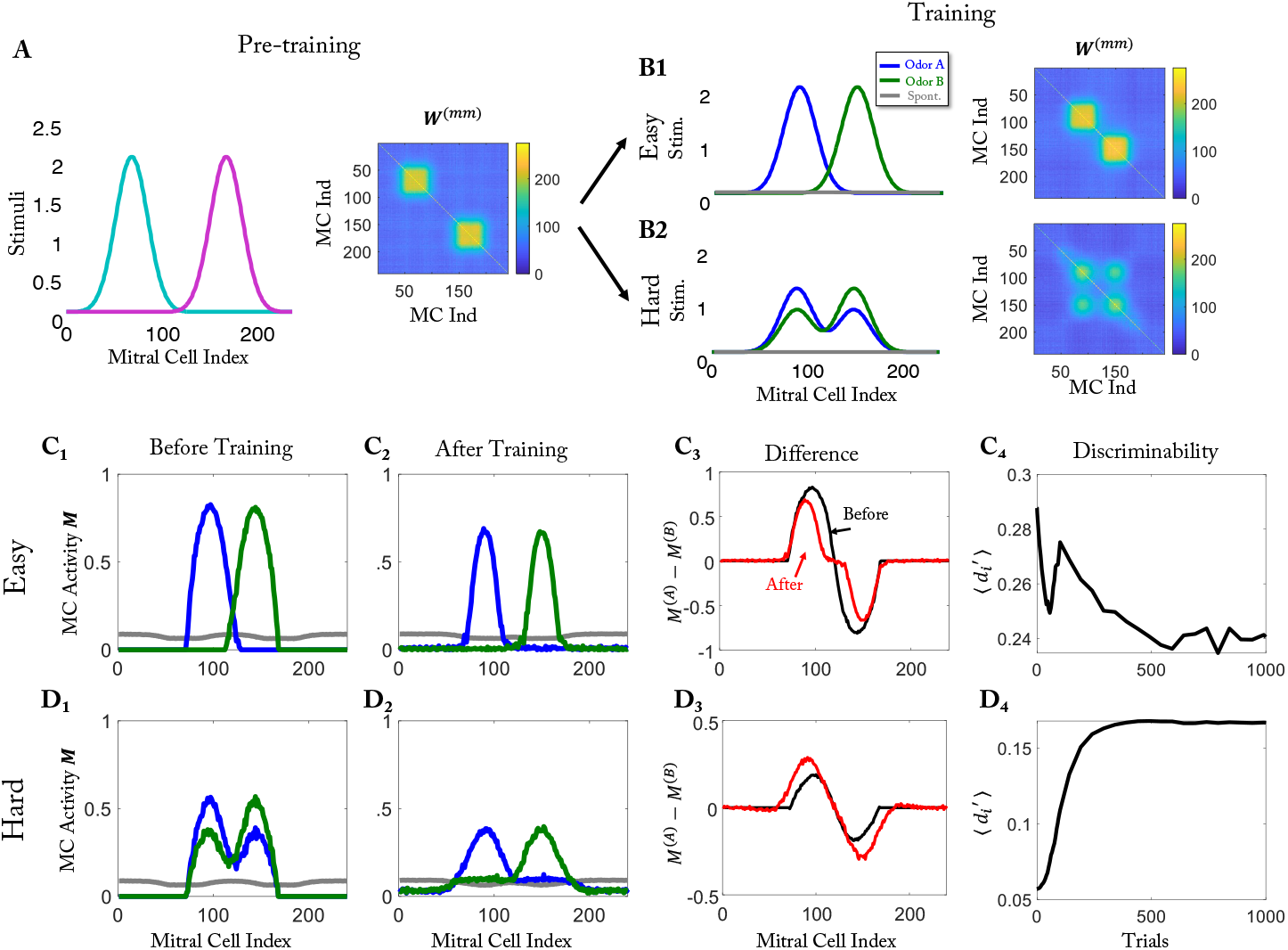
Visualization of the differential learning outcome for easy and hard tasks. (A) Stimuli for pre-training and effective connectivity matrix after pre-training. (B) Training stimuli for the easy task (B_1_) and the hard task (B_2_) and the resulting connectivity matrix after training. (C) Easy task: (C_1_) MC Activity before training (after pre-training). (C_2_) MC activity after training. (C_3_) Activity difference before (black) and after training (red). (C_4_) Temporal evolution of the discriminability during training. (D) As (C) but for hard task.

In the easy task, the effective connectivity resulting from the training had only two blocks, which were along the diagonal and corresponded separately to odors A and B (Fig 3B_1_). This inhibition reduced the MC activity and with it the number of responsive cells. Since most MCs responded essentially to only one of the two stimuli, the response difference decreased along with the overall response (Fig 3C_3_, red line vs. black line), decreasing the discriminability.

In the hard task, both stimuli activated the same set of MCs, but to a different degree, reflecting the difference in the concentrations of the two components. The discriminability of the stimuli was therefore not compromised if the MC activities were reduced, as long as the difference in the representations of the two stimuli was maintained. In fact, in that case the discriminability in terms of d’ was enhanced, since we associate reduced overall activity with reduced trail-to-trial variability (cf. Eq.(12)). The effective connectivity resulting from the training achieved this through the two additional, off-diagonal blocks (Fig 3B_2_). Each MC activated by component A received disynaptic inhibition from MCs that were activated by component A as well as from MCs activated by component B. With increasing concentration of component A the effective inhibition from the MCs activated by component A increased, while that from the MCs activated by component B decreased. Thus, the overall inhibition was relatively independent of the concentration of A and therefore about the same for both mixtures, preserving the response difference. In fact, for a large number of MCs the difference in the response even increased with training (Fig 3D_3_). Assuming that for a cell to be classified as divergent, this difference had to be above some threshold, the number of divergent cells decreased in the easy task, while it increased in the hard task (Fig 3C_4_,D_4_, cf. Fig S14C,D), as we found for the naturalistic stimuli (Fig 2D_2_,E_2_). Qualitatively similar results were obtained without pre-training phase (Fig S6).

### Odor specific adaptation

Repeated odor exposure was seen in [26] to lead to the response of a smaller number of MCs. A change in the response amplitude through repeated exposure has already been found earlier [25, 40]. Specifically, it was found that the repeated exposure to an odor over the course of a week substantially reduced the MC response specifically to that ‘familiar’ odor, but not to ‘novel’ odors, which the animal had experienced much less [25]. Such a differential response could be due to a modulation of the overall gain in the olfactory bulb by a signal that indicates the novelty or familiarity of the presented odors. Such a signal could arise some time after odor onset, since the odor has to be recognized as familiar first. A more parsimonious explanation is offered, however, by the activity-dependent spine dynamics of our plasticity model (Fig.4).

**Fig 4.**
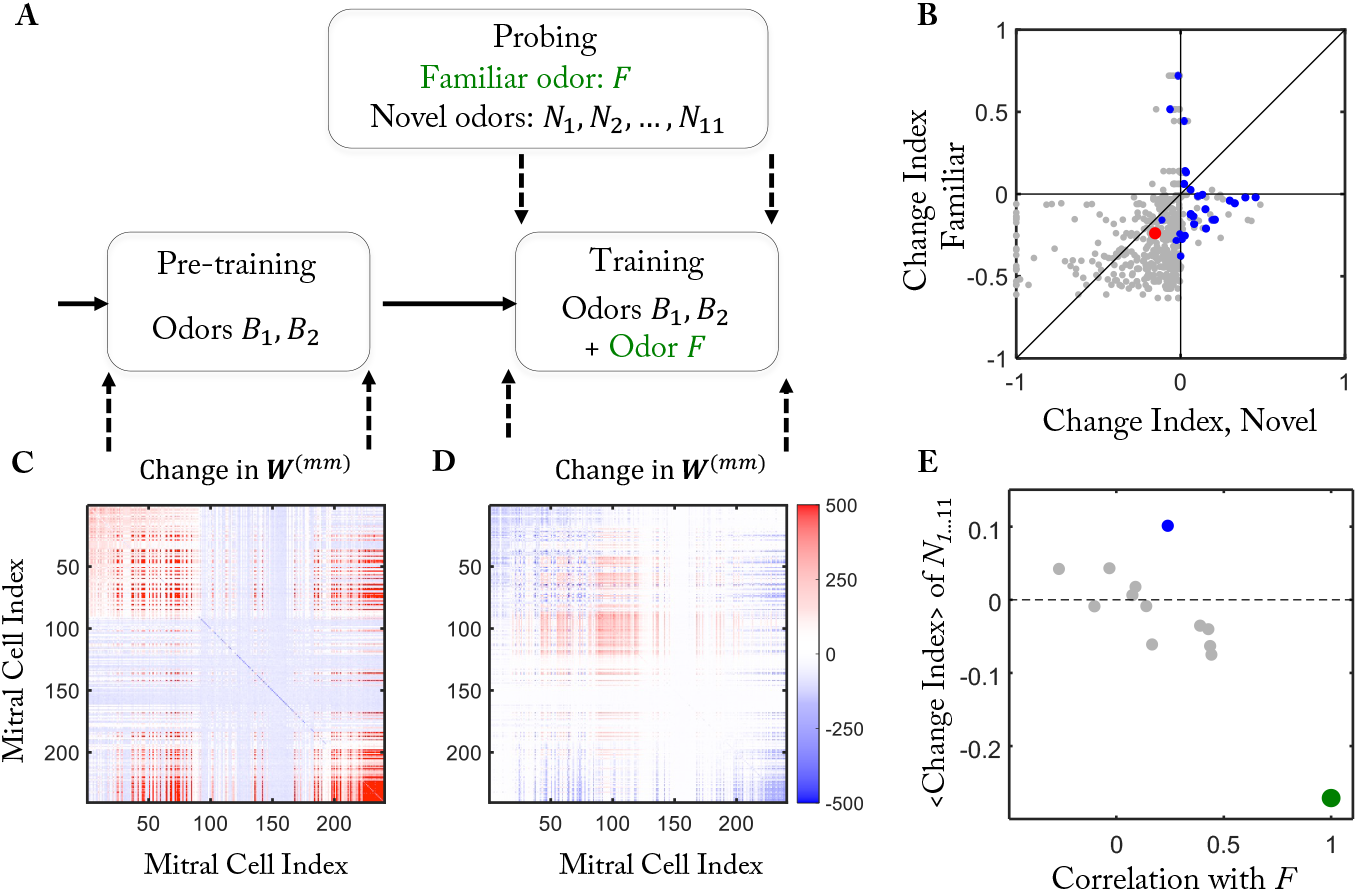
Odor specific adaptation (cf. [25]). (A) Simulation protocol. Pre-training with odors *B*_1,2_ established a background connectivity. During training the network became familiar with an additional odor *F*. (B) The MC response to the familiar odor *F* is more reduced than to the novel odors *N*_1…11_.). (C) During pre-training connections are predominantly added from GCs to MCs that are activated by the background odors *B*_1,2_ (MC indices below 90 and above 200). The color indicates the change in **W**^(*mm*)^, i.e. in the number of GCs connecting the respective MCs. (D) During training the number of connections of MCs responding to the familiar odor *F* increases, but that corresponding to odors *B*_1,2_ decreases (cf. (C)). (E) The mean change index of the novel odors *N*_1…11_ is correlated with their similarity to the familiar odor.

To mimic the procedure used in [25], we established a background connectivity by repeatedly exposing the network in the pre-training to two different naturalistic background odors *B*_1_ and *B*_2_ in an alternating fashion (Fig.4A). In the training period we included a third odor *F* in the odor ensemble as the ‘familiar’ odor and monitored the MC response to that odor as well as to 11 other, ‘novel’ odors *N*_1 11_. For each MC-odor pair we quantified the change in the network response using a change index. It was defined as the response of the MC to that odor at the end of the training period minus that at the end of the pre-training divided by the sum of those two activities [25]. As in [25], for the majority of MC-odor pairs the change index for the familiar odor was negative (Fig.4B). This reduction in response resulted from an increase in the number of GCs mediating the disynaptic inhibition among the MCs responding to the familiar odor (MC indices mostly between 30 and 150 in Fig.4D).

The network restructuring during the training generally also modified the MC responses to other, novel odors. For MCs that responded to a novel as well as to the familiar odor we compared the respective change indices. Since synapses that are not activated by the familiar odor do not become stabilized by the repeated odor exposure, the response of MCs to the novel odors was typically less reduced than that to the familiar odor, yielding a change index below the diagonal in Fig.4B, in agreement with [25].

The network restructuring during the training did not increase the inhibition for all MCs. For some MCs the inhibition was actually decreased leading to a positive change index (Fig.4B). Averaging across all responding MCs, the mean change index for a novel odor was negatively correlated with how similar it was to the familiar odor as measured in terms of the Pearson correlation of the stimuli (Fig.4E). A particularly strong reduction in the inhibition occurred among many of the MCs responding to novel odor *N*_4_ (marked blue in Fig.4B,D). This odor was actually identical to odor *B*_2_, which was used in the pre-training, where it established mutual inhibition among MCs with indices in the range 30 to 90 and above 200 (left panel of Fig.4C). During the training period, which included odor *F* in addition to *B*_2_, many of these connections were removed (Fig.4D), leading to a strikingly positive mean change index for that odor (blue circle in Fig.4E). Positive change indices for familiar and particularly for novel odors were also observed experimentally for quite a few MC-odor pairs in [25].

### Random spine dynamics can capture divergence and discrimination

So far we have seen that the model proposed here captured a variety of key experimental findings [25, 26]. A key prediction emerging from the activity-dependence of the spine dynamics is the formation of subnetworks of highly connected MCs and GCs that specifically respond to one of the training odors. We therefore asked whether the activity dependence is necessary to capture the experimental results and to what extent the experiments therefore allow to infer the formation of an odor-specific, learned network structure in the olfactory bulb. Alternatively, the spine dynamics could be statistically *homogeneous* with a density that is controlled by a global signal reflecting, e.g., a neuromodulator indicating the novelty or familiarity of an odor [41, 42].

As a control model, we implemented purely random, activity-independent spine dynamics, where the training with the familiar odor lead to a change in the mean number of MC-GC connections. Since the experiments [25] and the simulations (4) revealed a reduction in the MC-activity in response to the familiar odor, we increased during the training the mean number of connections between MCs and GCs, which is also consistent with an increase in the number of spines during perceptual learning [18].

For the hard task using simplified stimuli (cf. Fig.3) the fraction of responsive MCs decreased when the number *N*_*conn*_ of connections was increased, while the fraction of divergent MCs increased substantially over some range of *N*_*conn*_ (Fig.5B, cf. Fig.2). In parallel, the discriminability measured in terms of *d*′ increased (Fig.5C). This reflects the fact that increasing the random connectivity increased the inhibition for *all* MCs by roughly the same amount. In a linear firing-rate model this would preserve the difference between the MC responses; with the saturation of the MC response used in our model, the difference actually increased (Fig.5D,E).

**Fig 5.**
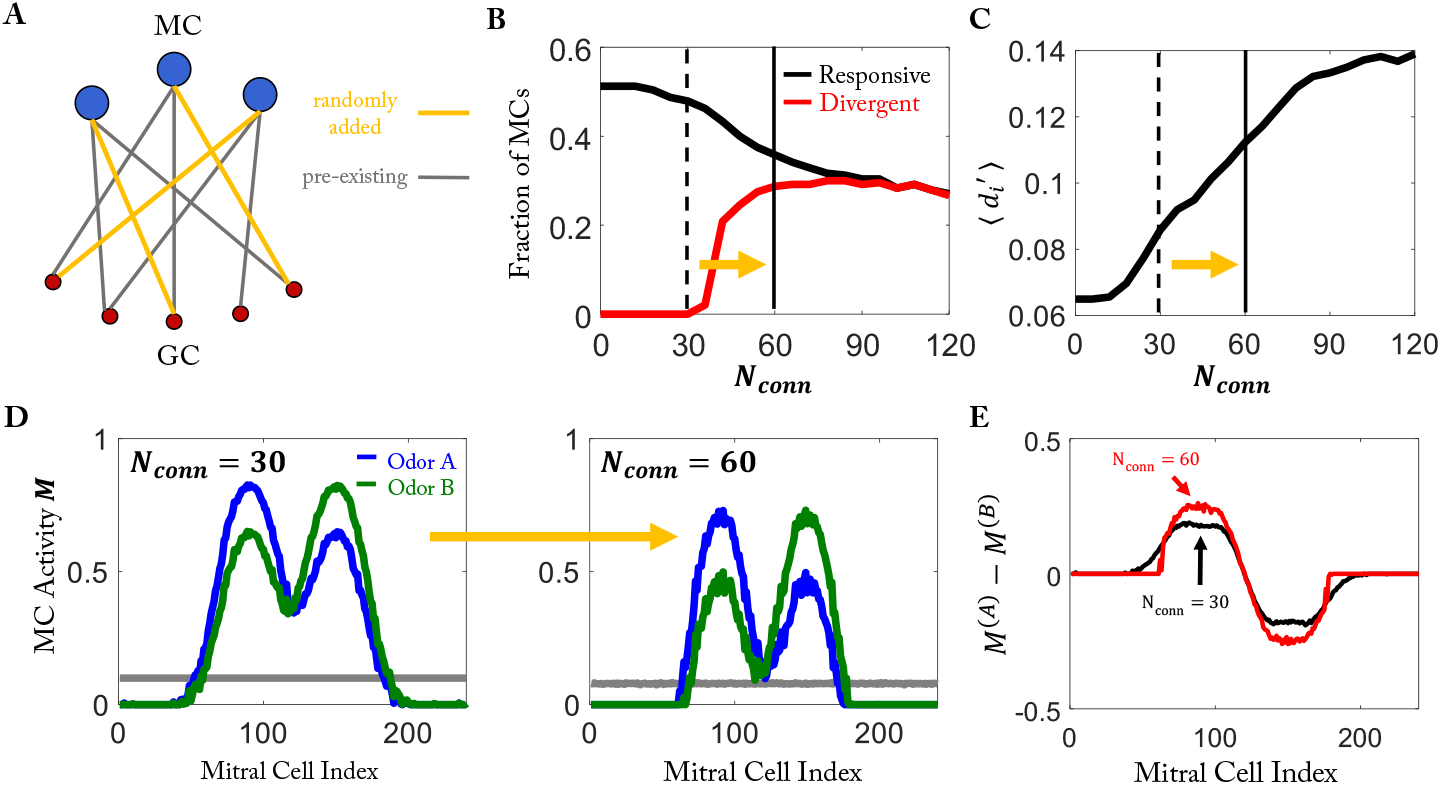
Random network model can capture enhanced differentiability. A) Random addition of connections during training. (B) Increased connectivity decreases fraction of responsive cells but can increase that of divergent cells (cf. Fig.2H). (C) Discriminability of the odors in the hard task increases with increasing *N*_*conn*_. (D,E) The difference in MC response does not decrease with increasing *N*_*conn*_ (cf. Fig.3I).

Thus, the aspects considered in Fig.5 are not able to distinguish between the activity-dependent and the random model. We therefore considered the results of the protocol of Fig.4 in more detail.

### Random spine dynamics fail to induce observed disinhibition

As in Fig.4, we exposed the random model in the pre-training to 2 background odors, added odor *F* as the ‘familiar’ odor to that ensemble during the training period, and probed the response to additional 11 ‘novel’ odors. As expected, and in contrast to the activity-dependent model, the random model showed no qualitative difference in the change index distribution of the familiar and the novel odors (Fig.6D). Strikingly, even though in the random model the mean change index for the familiar odor was less negative than that in the activity-dependent model (⟨*CI* ⟩_*random*_ = −0.14 vs ⟨*CI* ⟩ = −0.27), for none of the MC-odor pairs the change index was positive in the random model. In contrast, in the activity-dependent model about 8% of the MCs had a positive change index for the familiar odor (Fig.6E), which is in rough agreement with the experimental findings [25]. These MCs were disinhibited by the training (Fig.6A). For visualization purposes we also compared the changes in MC activity for the two models using simplified stimuli. Again, while in the random model the training led to a decrease in the activity of all MCs, in the activity-dependent model there were quite a few moderately activated MCs that were disinhibited by the training (Fig.6C).

**Fig 6.**
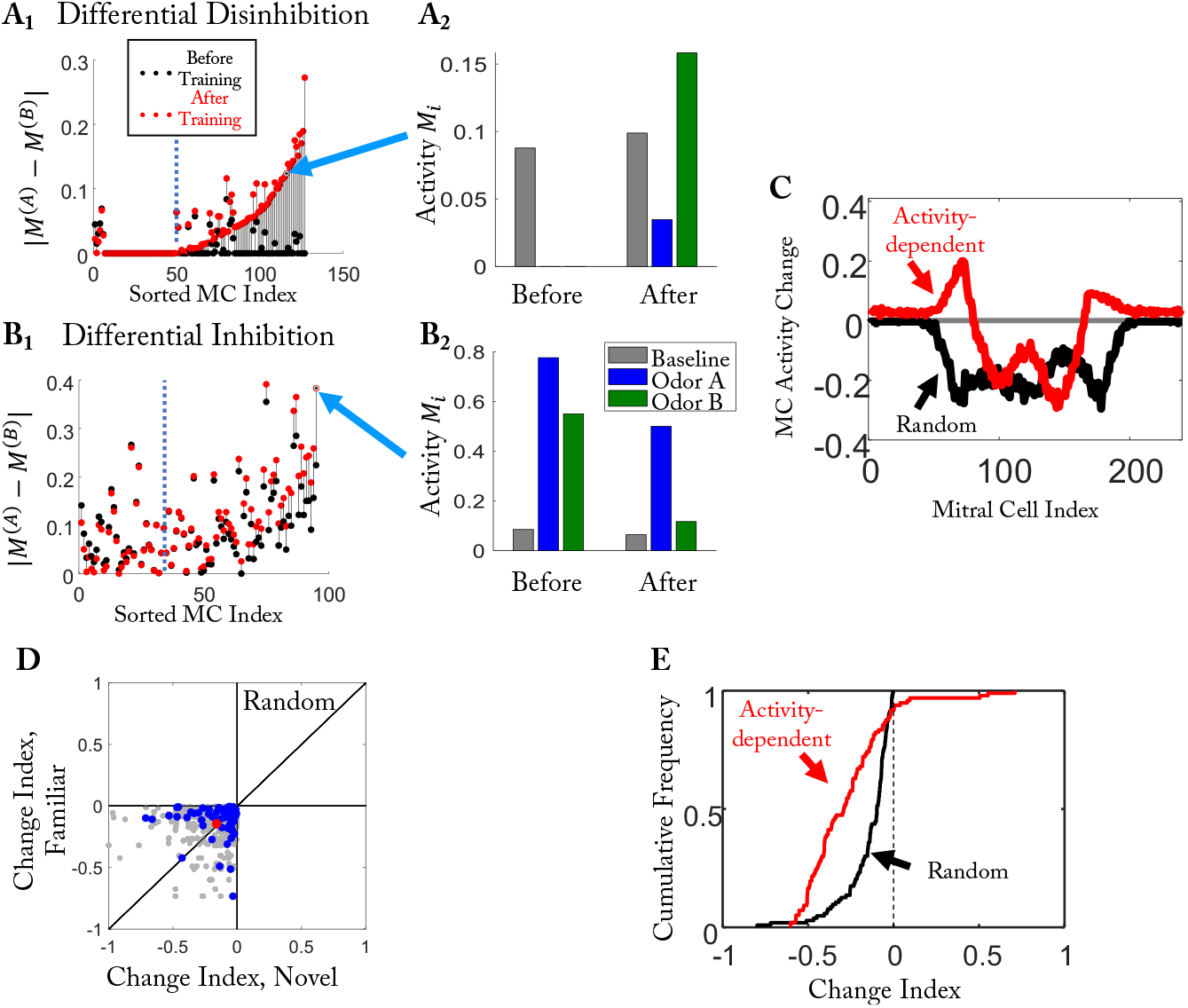
Disinhibition in the activity-dependent but not in the random model. (A) In the activity-dependent model training in the hard task with naturalistic stimuli (cf. Fig.2) disinhibited a large fraction of MCs differentially. The activity of a specific MC is shown in (A_2_). (B) In the activity-dependent model training also inhibited many MCs differentially. (C) Training-induced change in MC activities for the random and the activity-dependent model for the simplified stimuli. (D) In the random model the change index for familiar and novel naturalistic odors was statistically the same (cf. Fig.4B); Odor *N*_4_ is marked in blue. (E) The CDF of the change index shows disinhibition only for the activity-dependent, but not the random model.

A more detailed analysis of the results of the activity-dependent model showed that the disinhibition in the response of a given MC to two odors can be quite different even if the two odors are very similar to each other. Considering again the mixtures used in the hard task of Fig.2, we found a large number of MCs for which the difference in the response to the two stimuli was significantly increased by the network restructuring. For some of them the enhancement arose from differential disinhibition, wherein the MCs were inhibited by both odors before training, but only by one of them after training (Fig.6A). For other MCs the increased difference was due to differential inhibition. While these MCs were excited by both odors before training, they responded only to one of the very similar odors after training (Fig.6B). Both, differential disinhibition and inhibition were observed experimentally in [26] (their Supp. Fig.S1).

In contrast, in the random model no disinhibition was obtained. This is illustrated for one odor in Fig.6C. It also manifested itself in the change index for novel and familiar odors Fig.6D (cf. Fig.4B) and the cumulative distribution of the change indices (Fig.6E)

Thus, the improved discrimination of pairs of very similar stimuli and the reduced discrimination of dissimilar stimuli does not require the emerging connectivity to be odor-specific, but only an overall increase in inhibition. The observed disinhibition, however, is not consistent with random spine dynamics. But it is recapitulated with activity-dependent spine stability, which leads to odor-specific network structure. Thus, combined with the experimental observations of [26] our computational model supports the notion that odor exposure during perceptual learning leads to a network structure that reflects the presented odor.

### Removal of spines and forgetting by intermediate-amplitude re-activation

A key feature of our model is the non-monotonic behavior of the resilience function *ϕ* (Fig.1B), which renders spines stable not only if the associated GC is strongly active, but also when it is inactive. This allows previously learned connectivities to persist even if the corresponding odors are not presented any more. If MCs and GCs are weakly driven, however, the corresponding spines are removed. While the non-monotonicity of the resilience function is reminiscent of models for synaptic weight plasticity [34, 35], there is no explicit experimental evidence for it so far for the bulbar structural plasticity.

To illustrate the impact of the non-monotonicity of the resilience function we used simplified stimuli. During pre-training two dissimilar odors A,B were presented, which resulted in a connectivity comprised of two blocks on the diagonal (Fig.7A). In a subsequent training period we tested the persistence of the memory of the block corresponding to odor A when the network was re-exposed to that odor, but with a different amplitude (labeled *A*_1…4_ in Fig.7). When that amplitude was small, the connectivity remained essentially unchanged by the training (Fig.7B_1_). For intermediate activities, however, a fraction of the connections established in the pre-training were removed since the GC activity fell into the range in which the resilience function was negative (Fig.7B_2,3_). For large amplitudes the resulting GC activity was large enough to render the resilience function positive and the connections were stable (Fig.7B_4_).

**Fig 7.**
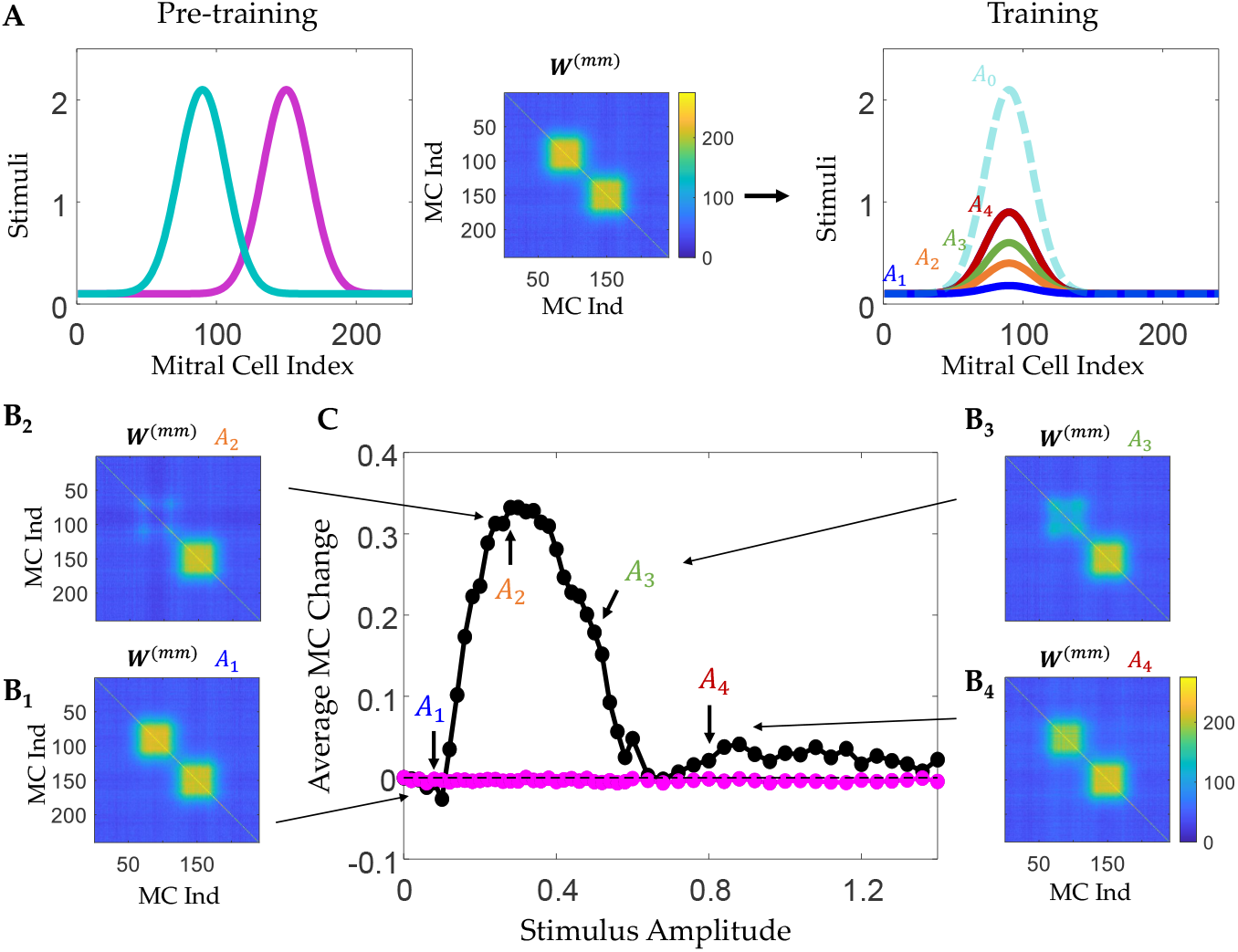
Removal of connections for intermediate GC activation. A) Pre-training with two odors, then training with only one of the two odors at different amplitudes *A*_1…4_. B) The connectivities **W**^(*mm*)^ resulting from the training depend non-monotonically on the stimulus amplitude *A*_1…4_ during training. C) Non-monotonic mean change in the response to odor *A* after training as a function of the amplitudes *A*_1…4_ used in the training.

Experimentally the connectivity itself is only poorly accessible. One can, however, probe the predicted changes using the response of the MCs to odor *A*, similar to the experiments in [25]. When the connections are retained during the training, the response of the MCs to that odor will not change compared to their response before the training. However, if intermediate stimulus amplitudes are used in the training, the removal of the connections manifests itself in an increase of the mean response of the MCs to odor *A* after the training (Fig.7C). Correspondingly, the distribution of the differences between the MC activities after and before the training shifts and broadens in a non-monotonic fashion when the amplitude of the stimulus used in the training is increased (Fig.S2).

Thus, if the odor exposure experiments in [25] were to be continued with the same odor but at a suitably reduced concentration, the model predicts that the MC response to that odor would increase again within days rather than undergoing the slow recovery over a period of 1-2 months that was observed in the absence of that odor [25].

The protocol used in Fig.7 does not have a simple behavioral read-out. Moreover, it relies on a change in the overall activity of the MCs. Since the input to the MCs is pre-processed in the glomerular layer, which is thought to provide an overall gain control of the input [43], changes in the odor concentrations may not translate directly into corresponding changes in MC activation and the ensuing change in the connectivity predicted by the model. We therefore considered also a slightly modified protocol using mixtures composed of two dissimilar odor components. To reduce the impact of the overall gain control, only the concentration of one of the two components was varied (Fig.8A). Exposing in pre-training 2 the network to two mixtures in which that component had similar but large values (Fig.8A_2_) enhanced the discrimination between these two mixtures (teal-magenta line in Fig.8G), However, due to pre-training 2 the network developed strong mutual inhibition between the MCs responding to the two mixture components (Fig.8E). This allowed the strong component in the training stimulus (Fig.8A_3_) to suppress the response to the weak component completely (Fig.8B). Subsequent training with the mixtures in which that component was weak, removed many of those connections (Fig.8A_3_,F) and largely reversed that suppression (Fig.8C). This enhanced the discriminability of those mixtures above its initial value without any inhibition (blue-green line in Fig.8G).

**Fig 8.**
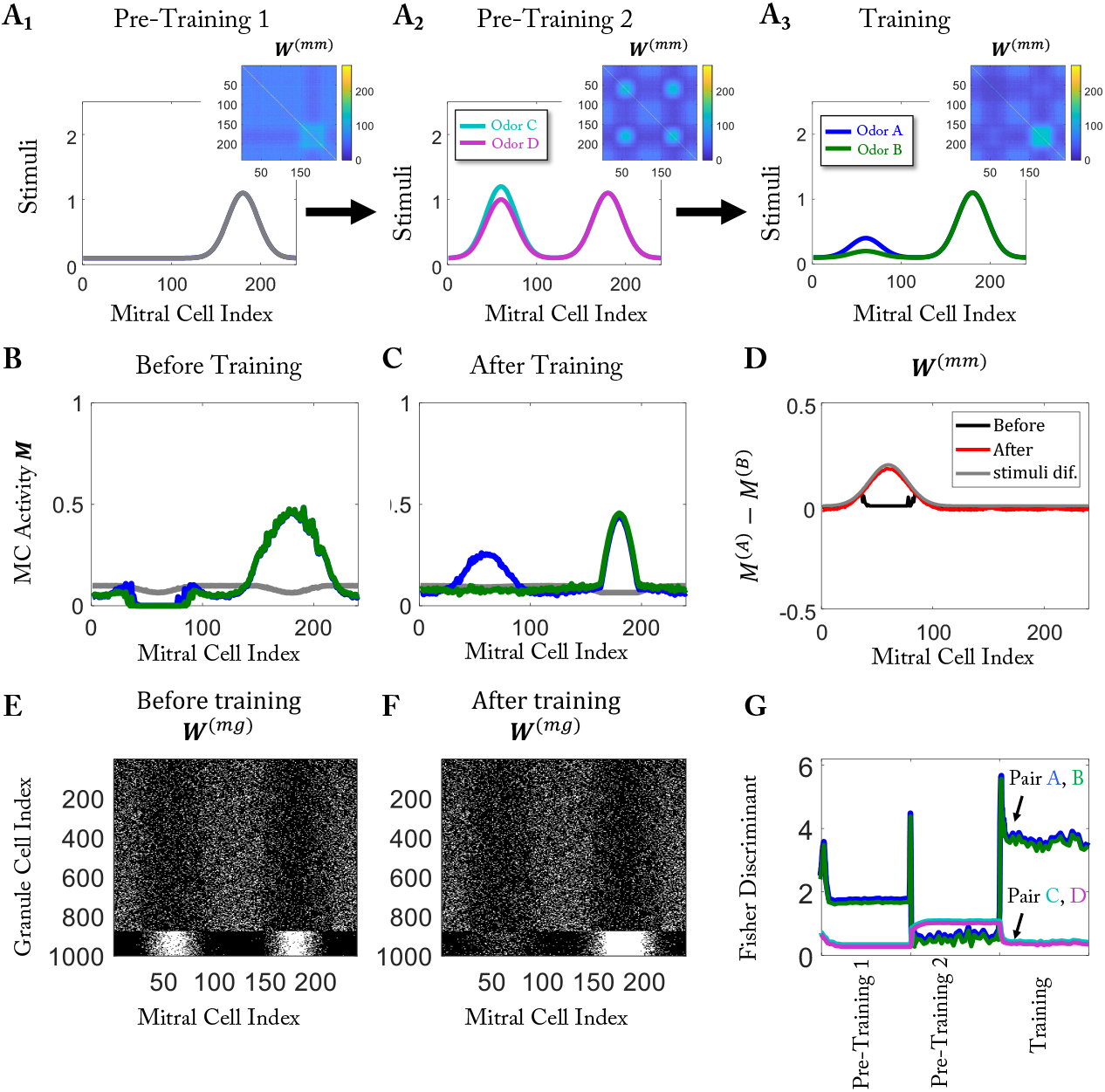
The discrimination of weak components of similar stimuli is abolished by training with strong components, but recovers through training with weak components. (A) Protocol: A_1_: pre-training 1 with only 1 odor, A_2_: pre-training 2 based on a mixture of that and a dissimilar odor, A_3_: training with a mixture in which the second component is only weak. (B) Response to the training stimulus before training; due to pre-training 2 the response to the weaker odor component is suppressed completely. (C) Training with the weaker component recovers that component. (D) MC response difference between odors A and B before and after training. (E,F) MC-GC connectivity before and after training. (G) Fisher discriminant for odor pair (A,B) and (C,D).

The removal of spines on GCs that are only intermediately activated lead also to forgetting of a discrimination task by learning a subsequent discrimination task, if the odors of the second task overlapped to some extent with those of the first task and therefore interfered with the retention of the relevant connections (Fig.S3). Upon re-exposure to the odors of the first task, the network very quickly re-learned to discriminate between those odors. In fact, the relevant connectivity was re-stablished faster than it had been learned in first odor exposure (Fig.S3G). In general re-learning occurred noticeably faster than the forgetting (Fig.S4).

### Realization of competition through competition for a limited resource

In order to keep a Hebbian model stable, different compensatory processes can be imposed depending on the state of nearby synapses on the same dendrite [37]. In the model that we have discussed so far we used top-*k* competition because of its simplicity, without identifying a specific, biologically feasible mechanism. In this section, we discuss one such mechanism.

The idea behind the competition is that within a single neuron, possibly within a single dendritic branch, resources used for the formation of synapses are limited. In hippocampal CA1 neurons, stimulating spines leads to the shrinking of neighboring unstimulated spines [44]. A plausible explanation of this phenomenon is that some resource is limited within the neuron. A computational model based on this idea has reproduced the fast multiplicative scaling behavior of synapses and explained a transient form of heterosynaptic plasticity [45]. Along similar lines, we assumed that the formation of each synapse depletes a resource pool by a fixed amount, and the formation rate is proportional to the fill-level of the resource pool. Further, we assumed that the resource for a synapse is returned to the pool when the synapse is removed and that the removal rate is independent of the resource pool size. We took the activation function (Fig 9A, red curve) to be comprised of a formation term, which depends on the fill-level of the resource pool (Fig 9A, blue curve), and a removal term, which is independent of the fill-level (Fig 9A, green curve).

**Fig 9.**
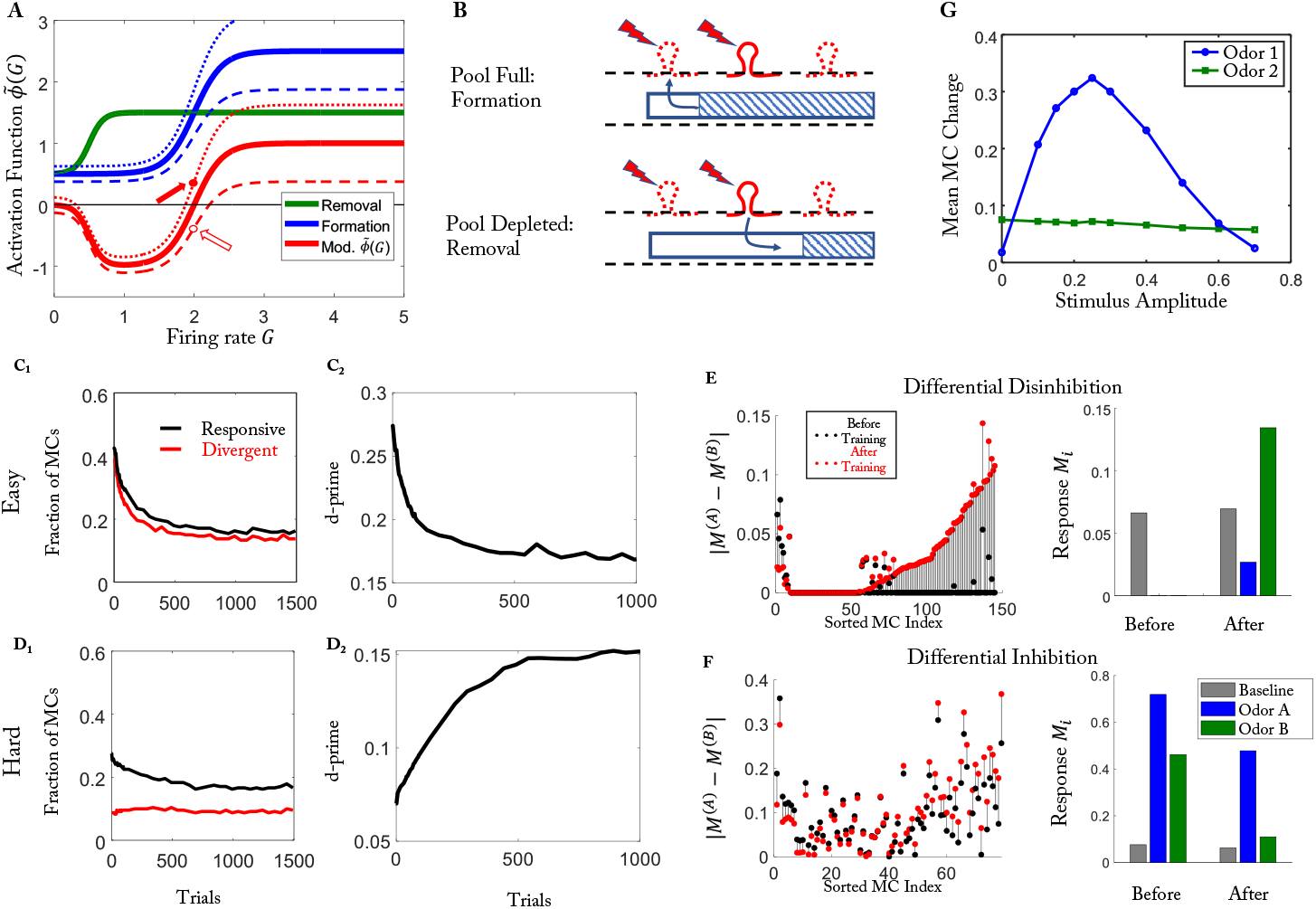
Synaptic competition via a common resource pool. The training protocol is the same protocol as in Fig 2. (A) The modified activation function 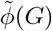 (red) is the difference between the formation function (blue) and the removal function (green). *P* = *P*_0_ (thick), *P > P*_0_ (dotted), *P < P*_0_ (dashed). (B) (Top) For *P > P*_0_ synapses tend to form (solid arrow in (A)). (Bottom) For *P < P*_0_ synapses tend to be removed even though the GC activity is the same (open arrow in (A)). (C1, 2) Responsive and divergent cells and discriminability for the easy task (cf. Fig.2E,F). (D_1,2_) Responsive and divergent cells and discriminability for the hard task (cf. Fig.2G,H). (E,F) Differential disinhibition and inhibition (cf. Fig.6). (G) Removal of spines for training with intermediate activities leads to an increase in MC activity (cf. Fig.7).

It is easy to see how the limited resource pool provides a homeostatic mechanism. For low pool levels synapses are removed (open arrow in (Fig 9A)), which replenishes the pool, while for high pool levels new synapses are formed (solid arrow in Fig 9A)), decreasing the pool level (Fig 9B). The equilibrium pool level and with it the number of synapses in the steady state is regulated by the activity of the GCs, which in turn depends on the number of synapses *via* the activity of the MCs connected to the GCs.

With this biophysically plausible stabilization of the structural plasticity, our model yielded results that were qualitatively similar to those obtained with the simpler, top-*k* competition model (compare Fig 9C-F with Fig 2D,E and Fig S5 with Fig 3).

## Discussion

In the rodent olfactory bulb structural plasticity in the form of the extensive formation and removal of synaptic spines persists into adulthood [6] and may be a key factor in establishing and maintaining the network structure of the olfactory bulb. Since it is operating on timescales that are comparable to some learning processes [6, 18] it may play an important role in these processes. In this study, we proposed a simple Hebbian-type model to investigate this role. Our model shows that a local unsupervised learning rule according to which the formation and removal of spines depends on GC and MC activity is consistent with and explains a host of experimental observations [18, 25, 26]. Among them is the intriguing observation that training can have opposite impact on the discriminability of very similar and of dissimilar odor pairs. As observed experimentally, in our model training enhanced the discriminability of similar odors, but it reduced the discriminability of the training odors if they were quite different from each other. In fact, for the dissimilar odors, the number of MCs that responded differently to the odors decreased with learning, deteriorating the performance.

Since a key prediction of our activity-dependent model is that upon repeated exposure with relevant odors the olfactory bulb develops subnetworks associated with these odors, we investigated whether the experimental results are also consistent with a model in which learning modifies the inhibition in a random, statistically homogeneous fashion, independent of the activity, and therefore without forming odor-specific subnetworks. Surprisingly, the random model also captured the differential outcome of training with similar and with dissimilar odor pairs. It failed, however, to recapitulate the observation that during that learning process, which led to an overall increase in the number of inhibitory connections, a sizable fraction of the MCs became more, rather than less active, i.e. they were disinhibited rather than inhibited [26]. Similarly, the activity-dependent, but not the random model, captured the odor-specific reduction in MC activity that is observed after an animal has been familiarized with an odor, which is much more pronounced for that familiar odor than for less experienced odors [25]. One could imagine that the reduced response to a familiar odor is the result of an inhibition-enhancing global modulatory signal that may be triggered by the recognition of the odor as familiar. This would, however, not explain the significant fraction of MCs that are disinhibited in response to the familiar or the novel odor [25]. Thus, in combination with [25, 26] our model provides strong indirect evidence that perceptual learning leads to the formation of MC-GC subnetworks comprised of MCs and GCs having significantly overlapping receptive fields.

Moreover, the model offers a foundation for further studies of learning-related functions in the olfactory bulb. For instance, in the current version of the model the novelty or salience of an odor is not considered explicitly. Only implicitly it is assumed that the presented odors have sufficient salience, since experiments suggest that the plastic changes occur predominantly for novel or otherwise salient odors [19, 46].

In contrast to a previous model for structural spine plasticity [6], the model presented here describes a learning mechanism that allows the network to remember previously learned tasks in its connectivity. These memories are robust against subsequent learning of dissimilar tasks. The learned connectivity is only erased by exposure to interfering stimuli, which overlap significantly with previously learned ones. This prediction should be amenable to experimental tests using sequential perceptual learning [47] with two odor pairs, each comprised of two similar odors. Repeated exposure with one of the two odors of the first pair will enhance the spontaneous discriminability of the odors in that pair. The subsequent repeated exposure with an odor from the second pair is expected not to affect the discriminability of the odors in the first pair, if the odors in the second pair are dissimilar from the first pair. However, if there is a significant overlap between the pairs, the second training is expected to compromise the discriminability of the first pair. Note, however, that if the odors in the second pair are very similar to those in the first pair, this interference is not expected. The latter aspect would differentiate this forgetting from the widely observed retrieval-introduced forgetting [48].

When assessing the role of structural spine plasticity in forgetting, attention has to be paid to the time scales of the experiments. In recent experiments the successive perceptual learning and forgetting of different odor pairs has been investigated [4]. There it was found that learning a second pair of odors seven days after the training with the first pair strongly interfered with the memory of the first pair, while learning after 17 days did not. The experiments indicated that this interference arose from the enhanced apoptosis of adult-born GCs that was triggered if the time between the training periods was only seven days. Since the results of [25] suggest that significant changes in the connectivity arise already within the first day of exposure, performing the interference experiments on that shorter time scale may avoid complications arising from the apoptosis found in [4] and may focus on the role of the structural spine plasticity described by our model.

We discussed two fast homeostatic mechanisms that stabilize the network activity despite the Hebbian-type learning rule: an abstract top-*k* competition and a biologically plausible competition of synapses for a limited resource that operates via a sliding threshold for the formation of synapses, somewhat reminiscent of the BCM-model [34]. The overall behavior of both models did not depend much on which homeostatic mechanism was at work. It is worth noting that the specifics of the stabilization mechanism may vary across brain areas. Moreover, resource-dependent competition may be too local to lead to competition across different dendrites or the whole neuron. In addition, heterosynaptic plasticity may be controlled more by activity rather than a limited resource [44]. For these reasons, we leave open the question of how to implement the stabilization mechanism.

In principle, activity-dependent network restructuring could also arise from synaptic weight plasticity rather than structural plasticity. Recent experiments show, however, that in adult-born GCs the amplitudes of evoked IPSCs were not significantly changed after learning [18]. This suggests that changes in the inhibitory synaptic weights are not likely to be the dominant mechanism for learning. At the same time, in the same experiment, the frequency of the evoked IPSCs was significantly up-regulated after learning, which highly suggests an increase in the number of synaptic spines [18].

What aspects of olfaction might favor structural plasticity even in adult animals, despite its seemingly high metabolic cost? A characteristic feature of olfaction is the high dimensionality of odor space. As a consequence, the layout of the olfactory bulb does not allow substantial chemotopy and MC dendrites have to reach across large portions of the bulb [28, 49], where they establish only a very sparse connectivity with GCs. To wit, each of the about 5 million GCs connects with on the order of 50-100 of the 50,000 MCs [50]. Thus, only less than 1% of the possible connections are actually made. The sparseness of the connectivity has recently been identified as a key factor favoring structural plasticity over synaptic-weight plasticity in the context of feedforward networks [51]. This may also be the reason why structural plasticity seems to be essential in the adult motor system, as has been found recently [52].

A characteristic feature resulting from the structural plasticity implemented in our model is the emergence of MC-GC subnetworks that are specific to the learned stimuli. The interaction between the excitatory MCs and the inhibitory GCs is known to drive the *γ*-rhythms that are prominent in the olfactory bulb [53]. Since their power is enhanced with task demand [54] and since the optogenetic excitation of GCs in the *γ*-range enhances discrimination learning [55], they are presumed to play a role in odor processing. The different subnetworks that are predicted by our model to emerge from learning different odors are likely to generate each its own *γ*-rhythm with its own frequency. The question then arises whether these rhythms will synchronize if the bulbar network is driven by the corresponding odor mixtures. Depending on the temporal window over which the bulbar output is read out in downstream brain areas [56, 57], the perception of such mixtures may then well vary with the degree of synchrony of the different rhythms. Interestingly, the synchrony of such interacting rhythms can be enhanced by uncorrelated noise and by neuronal heterogeneity *via* a paradoxical phase response [58, 59].

A prominent feature of the olfactory bulb is its extensive centrifugal input *via* top-down projections that target particularly the GCs. Our model for the structural spine plasticity suggests that the GC develop receptive fields for specific odors of the training set and are preferentially connected with MCs with similar receptive fields. Due to the reciprocal nature of the MC-GC synapses, this implies that the activation of such a GC inhibits specifically MCs with the same response profile. If the top-down connectivity was able to target specific GCs, top-down inputs could modulate bulbar processing in a controlled fashion by inhibiting specific MCs. Computational modeling [15] suggests that the extensive adult neurogenesis of GCs observed in the bulb naturally leads to a network structure that allows just that. That model predicts that non-olfactory contexts that have been associated with a familiar odor allow to suppress that odor, enhancing the detection and discrimination of novel odors. The spine plasticity investigated here leads to a network structure that is similar to the bulbar component of the network obtained in the bulbar-cortical neurogenesis model. Structural spine plasticity is therefore expected to support the specific control of bulbar processing by higher brain areas.

## Methods

The computational model comprises MCs and GCs (Fig 1A) that are described by the firing-rate models

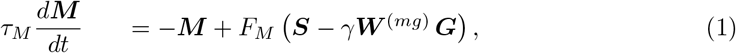

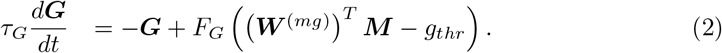

Here the vectors ***M*** and ***G*** represent the firing rates of *N*_*MC*_ MCs and *N*_*GC*_ GCs, respectively. The MCs receive sensory inputs ***S*** from the glomerular layer and inhibitory input from GCs, with *γ* denoting the inhibitory strength. The GCs receive excitation from MCs, with *g*_*thr*_ *>* 0 setting a non-zero firing threshold for the GCs. Reflecting the reciprocal nature of the MC-GC synapses the connectivity matrix ***W*** ^(*gm*)^ is the transpose of ***W*** ^(*mg*)^. Focusing on the structural plasticity of the synapses, we do not include synaptic weight plasticity, the characterization of which is still somewhat incomplete [31, 32]. We therefore keep the weight *γ* fixed and set ***W*** ^(*mg*)^ either to 1 or 0. Initially, each row of ***W*** ^(*mg*)^ has *N*_*conn*_ non-zero entries, reflecting that each granule cell connects to *N*_*conn*_ mitral cells.

Reflecting the large spread of the secondary dendrites of MCs [28], we allow synapses to be formed between all MCs and all GCs without any spatial limitations. Thus, the ordering of the MCs and the GCs is arbitrary.

The firing rates depend on the neurons’ inputs via the activation functions *F*_*M*_ and *F*_*G*_ given in (3,4), which are taken to be sigmoidal and piecewise linear, respectively. MCs within the olfactory bulb experience saturation behavior. In [60] the authors recorded responses of MCs when the concentration of the odorant changed 10-fold. Among the recorded cells, 38% of MCs responded linearly to stimuli, and 29% of the MCs experienced saturation behavior. This saturation may be the result of saturation in the inputs to the MCs or in the MCs themselves. The activity of the sensory neurons saturates on a log scale when the odor concentration changes 1000-fold [61]. For the concentration changes (0.6 : 0.4 mixture or pure odorant) relevant in our studies, we therefore assumed that saturation occurred mostly in the MC rather than their sensory inputs and took the activation functions in (1,2) to be

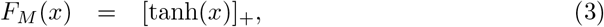

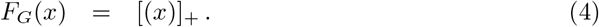

Based on the observations in [5, 22, 23, 33], we assume the rate of formation and removal of a synapse connecting MC *i* and GC *j* to depend on the activities of those neurons. As a proxy for those activities we take the steady-state solutions of Eqs.(1,2) for given stimulus **S**. We express the formation and removal rates in terms of a single rate function,

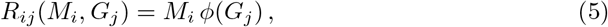

where *M*_*i*_ and *G*_*j*_ are the respective steady-state activities. The activation function *ϕ*(*G*_*j*_),

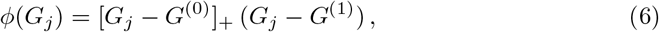

changes sign at a threshold *G*^(1)^, which controls whether a new synapse is formed or an existing synapse is removed (Fig 1B), giving the structural plasticity a bidirectional dependence on the GC activity. In addition, there is a second threshold, *G*^(0)^, below which the structural change is negligible. Specifically, in each trial of duration Δ*t*, which is one step in our simulation, a new synapse is formed with probability 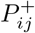,

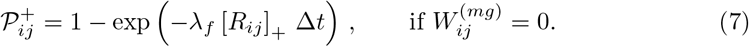

Conversely, an existing synapse is removed with probability 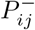

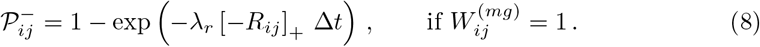

Here, *λ*_*f*_ and *λ*_*r*_ are the formation and removal rates, respectively.

Since the change in the total number of synapses appears to be limited in experiments [36, 62], we introduce a homeostatic mechanism that limits the maximal number of synapses on a given GC. For most of our results, we employ a top-*k* (*k > N*_*conn*_) competition mechanism, in which only the *k* synapses with the largest values of *R*_*ij*_ survive if a GC has more than *k* synapses. A soft top-*k* competition can be realized by a resource-pool competition with additional assumptions, as discussed in the section Realization of competition through competition for a limited resource. However, other mechanisms like scaling [63] or the ABS (Artola, Bröcher and Singer) rule [64] may alternatively restrict the total number of synapses. Since the stability mechanism was not our focus, we mostly used the (hard) top-*k* competition without directly specifying its biological cause.

Summarizing the algorithm, in each step the steady-state firing rates ***M*** and ***G*** in response to a randomly chosen odor ***S*** from the training set are computed. Based on the resulting formation/removal rates *R*_*ij*_ the connectivity matrix ***W*** ^(*mg*)^ is updated in two parts. First, we implement homeostasis by top-*k* competition: if any GC has more than *k* synapses, the corresponding number with the smallest values of *R*_*ij*_ are removed. Then, in the learning step the connectivity is updated based on the Hebbian learning rules (7, 8), except that synapses that were just removed in the homeostasis step are not recreated in this time step. We simulate the model long enough for the connectivity to reach a statistically steady state as measured by the discriminability of the training odors (see Sec.Characterizing Discriminability). As initial condition we take a connectivity in which each GC is randomly connected to *N*_*conn*_ MCs. Unless specified otherwise, the model parameters are as listed in Table 1. The steady states are obtained by setting *τ*_*G*_ = 0, solving Eq.(1) using ODE45 in Matlab, with ***G*** solved from Eq.(2). We found that the overall behavior of the model is robust with respect to changes in its parameters (Supplementary Materials).

Instead of the hard top-*k* competition the number of synapses can also be limited by a resource-pool competition. To this end we replace the activation function *ϕ*(*G*) in (5) by a function 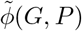 that also depends on the fill-level *P* of that resource pool (Fig 9A, red curve) and assume that it is comprised of a formation term 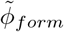 (Fig 9A, blue curve) and a removal term 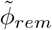 (Fig 9A, green curve),

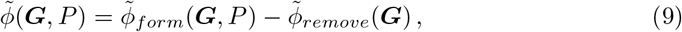

which are, respectively, given by

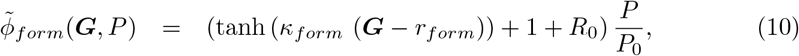

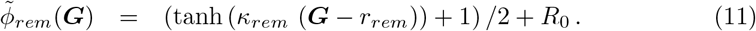

The fill-level *P* of the resource pool depends on the current number of synapses *n, P* = *P* ^(*all*)^ - *n*, where *P* ^(*all*)^ is the total amount of the resource in the cell, which was assumed to have the same constant value for all GCs. Here *P*_0_ is the equilibrium size of the resource pool for which no spines are formed or removed when the GC is not active. Thus, for low GC-activity *P* eventually goes to *P*_0_ and the number of the synapses on that GC goes to the initial value *n* = *N*_*conn*_ = *P* ^(*all*)^ - *P*_0_.

In analogy to the original model, we define the two thresholds 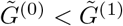 *via* 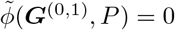. The formation of synapses on significantly active GC is controlled by 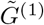. As more synapses are formed and the fill-level decreases, 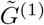 increases. This makes it more difficult for synapses to form on that GC, leading to a saturation of the number of its synapses. This dependence of the threshold 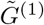 is reminiscent of the sliding threshold of the BCM model for synaptic-weight plasticity [34]. However, the sliding is caused here by the changes in the limited resource instead of a temporal average over the previous activities. The change in 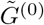 is opposite to the change in 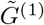 when the size of the resource pool changes.

In [26], mice were exposed to the test odors only when they were on the training stage. When they were back in the cage, the training odors were absent, and the activities of GCs corresponding to those odors were likely to be low. In the original model, the activation function 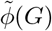 vanishes when *G < G*^(0)^, which means synapses are neither formed nor removed. Thus, the network is static. However, in the resource-pool model the activation function 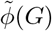 is in general non-zero when *G < G*^(0)^. As a result, periods of low input also lead to changes in the connectivity. We include such ‘air trials’, during which only spontaneous activity is presented, after every four training trials. During air trials, the GC activity is very low and synapses are formed and removed based particularly on the size of the respective resource pool, pushing each pool towards *P*_0_. Given the uniform input to the MCs during air trials and the dependence of spine formation on MC activity (cf. (5)), the formation is biased towards MCs that receive less inhibition, i.e., that have fewer connections with GCs, while the removal is biased towards MCs that receive more inhibition. We find that inserting such air trials increases the dynamics of the synapses and speeds up the convergence of the network evolution. It may compensate for the fact that in the resource-pool model an existing synapse is removed only if the activation function is negative, 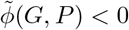, while in the top-*k* competition model, synapses can be removed not only when *ϕ*(*G*) *<* 0, but also directly by that competition.

The top-*k* mechanism and the competition *via* a resource pool lead to similar results. We expect that this would also be the case for other mechanisms that may restrict the total number of synapses, like scaling [63] or the ABS (Artola, Bröcher and Singer) rule [64]. Since the stability mechanism was not our focus, we mostly used the (hard) top-*k* competition without directly specifying its biological cause.

The additional parameters of the pool-competition model are listed in Table 2.

**Table 2.** Table of parameters for the resource-pool model (unless stated otherwise).

|  |  |  |  |
| --- | --- | --- | --- |
| $\kappa_{form}$ | 2.5 | $\kappa_{rem}$ | 5 |
| $r_{form}$ | 2 | $r_{rem}$ | 1 |
| $R_0$ | 0.8 | $P_0$ | 20 |
| $\Delta t$ | 1 | $\gamma$ | $4.2 \cdot 10^{-4}$ |
| $\lambda_f$ | 1 | $\lambda_r$ | 1 |

### Generation of simplified stimuli and naturalistic stimuli

The naturalistic stimuli are based on glomerular activation data [39], which contains 2-*d* imaging z-scores of the glomeruli responses for a variety of odorants. In our simulations, we used carvone, citronellol (pre-training), ethylbenzene, heptanal (training, pure, and mixture). For the novel odors employed in Figs.4,6 we used in addition limonene, ethylvalerate, 2-heptanone, acetophenone, valeric acid, isoamylacetate, isoeugenol, 1-pentanol, p-anisaldehyde. Since in the original data, not all pixel values were available for all odorants, we kept only those that were common to all of them. We then downsampled the resulting 2074 data points to 240 sample points ***S***^(*orig*)^.

The z-score data ***S***^(*orig*)^ do not include the baseline activity of the glomeruli. Thus, a z-core of 0, which represents the mean activity across the whole population, does not imply that the activity was not affected by the odor presentation. To obtain a rough calibration of the mean activity, we used the observation that around 60% of the MCs are activated by any given strong stimulus [25]. In the absence of other information, we used this as a guide to re-calibrate the 40% percentile z-score *S*^(40%)^ as 0. Then, we normalized the data by their maximum, resulting in

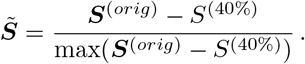

Further, MCs are activated by the airflow even without any odorants, and odor representations change mostly linearly with the concentration of odorants [60]. For any given mixture with concentration *p* of odor *A* and (1 - *p*) of odor *B* we therefore modeled the stimulus as

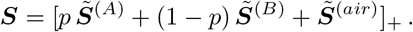

Here 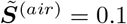 is the ‘air’ stimulus and [·]_+_ the rectifier, [*x*]_+_ = max(*x*, 0).

For the simplified model stimuli we used scaled Gaussian functions with different mean and standard deviation as 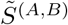 and set 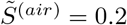.

### Definition of responsive cells and divergent cells

We defined the response of an MC as the change in activity induced by an odor compared to the activity with air as stimulus. Specifically, if a MC had activity ***M*** ^(*A,B*)^ in response to stimuli ***S***^(*A,B*)^, and it had activity ***M*** ^(*air*)^ when presenting air as a stimulus ***S***^(*air*)^, we defined the response to odor *A, B* as 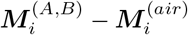.

The definitions of responsive and divergent cells were adapted from [26]. A cell was classified as responsive, if its response to one odor in an odor pair was above a given threshold *θ*, i.e. 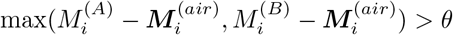. A cell was classified as divergent, if the difference in its response to the two odors was above the same threshold, 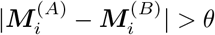. Our results were robust over a range of values of the threshold *θ* (Fig S14).

### Characterizing Discriminability

We used d-prime and the Fisher discriminant to assess the discriminability of activity patterns. For each divergent neuron d-prime is given by the difference between the mean activities of the neuron for the two presented odors scaled by the combined variability of the activities. In our firing-rate model (1,2) the MC-activity exhibits fluctuations only on the time scale of network restructuring, which is much longer than the odor presentation. Assuming, as in [15], that the MC firing rates 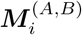 represent the mean values of independent Poisson spike trains, the variances **Σ**^(*A, B*)^ of those spike trains are given by their respective means, 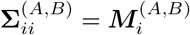 and 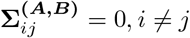. Thus, for each MC we have

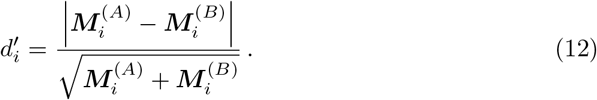

One way to characterize the overall discriminability of the activity patterns is then given by the average 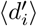 of the d-prime values across all neurons, as was done in [26]. This average 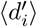 may decrease when additional, less-discriminating MCs are included in the ensemble. However, the information transmitted by the ensemble does not decrease. We therefore used also the Fisher discriminant to measure the discriminability. In general, it is given by

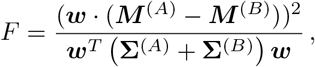

where ***w*** is an arbitrary vector, which can be interpreted as the weights with which the MC activity is fed into a linear read-out neuron. The weights ***w*** that maximize *F* are given by **w**^(*opt*)^ = (**Σ**^(*A*)^ + **Σ**^(*B*)^)^*-*1^(***M*** ^(*A*)^ - ***M*** ^(*B*)^). The optimal Fisher discriminant used to assess discriminability is then given by

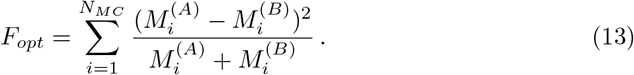

It is related to the d-prime values of the individual neurons *via*

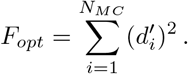

Thus, *F*_*opt*_ always increases with the addition of MCs, reflecting the fact that even poorly discriminating MCs provide some additional information about the odors. The optimal read-out **w**^(*opt*)^ puts less weight on the poorly discriminating MCs to avoid that the trial-to-trial variability associated with those MCs deteriorates the overall discrimination.

## Acknowledgments

We thank S. Saha, K.A. Sailor, and P.-M. Lledo for discussions and for sharing their data with us prior to publication. We gratefully acknowledge discussions with T. Komiyama.

This work was supported by funding from the National Institutes of Health (https://www.nidcd.nih.gov/) through grant DC015137 to HR.

## Data availability

The authors declare that the data supporting the findings of this study are available within the paper and its Supplementary Information files or from the corresponding author on reasonable request. A sample code is available in the Supplementary Information files.

## Author information

### Contributions

J.M. and H.R. conceived the ideas of this study and designed the model. J.M. carried out the simulation and analyzed the data. Both contributed to the writing of the manuscript.

### Corresponding authors

Correspondence to Hermann Riecke.

## Supplementary Materials

### Randomized Connectivity

To further illustrate that the network learned not by simply changing the number of synapses, but by developing a specific, stimulus-dependent connectivity, we assessed the performance of the network when the connections of each GC were rewired to random MCs with probability *p*_*rewire*_ while keeping the number of synapses on each GC the same. For *p*_*rewire*_ = 0 (Fig S1A), the connection was the originally learned network; for *p*_*rewire*_ = 1, the connection for each GC was totally random (Fig S1C). Indeed, for the highly similar training odors, the discriminability predominantly decreased as the randomness is increased (Fig S1D). For the network trained on the dissimilar odors (easy task), the discriminability increased, as expected, when the randomness was increased (Fig S1F). Interestingly, when *P*_*rewire*_ was increased beyond ≈ 0.7 these trends reversed (Fig S1D, F) somewhat.

**Fig S1.**
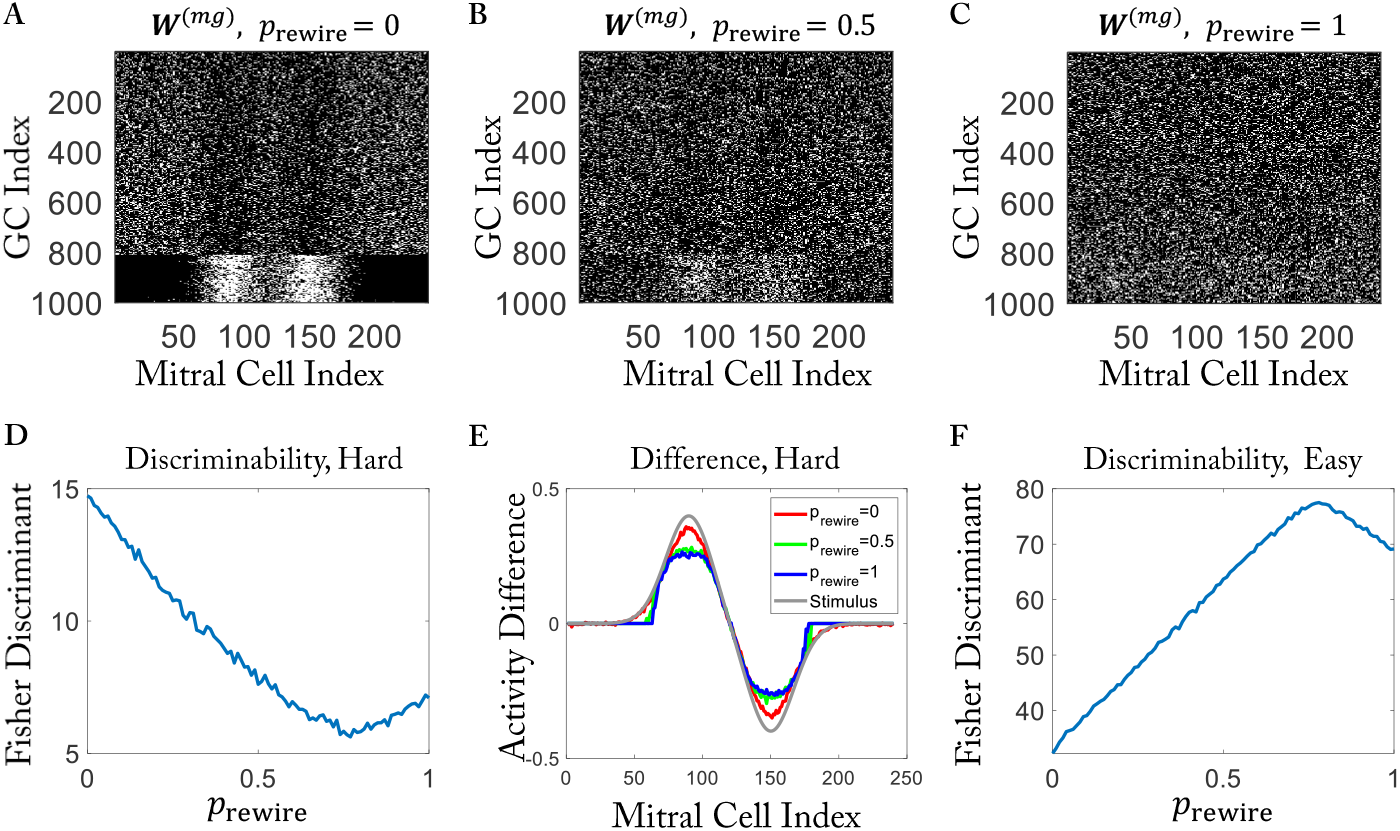
Randomly rewiring trained networks. (A to C) Connectivity matrices after rewiring a network that was trained with the hard task (Fig 3C, bottom). Rewiring probability *p*_*rewire*_ = 0, 0.5, 1, respectively. (D) For the hard task discriminability predominantly decreases with increasing rewiring probability *p*_*rewire*_. (E) Activity differences are reduced after rewiring. (F) For the easy task discriminability predominantly increases with increasing *p*_*rewire*_ (cf. Fig 3C, top).

### Removal of Spines for Intermediate GC Activation (cf. Fig.7)

**Fig S2.**
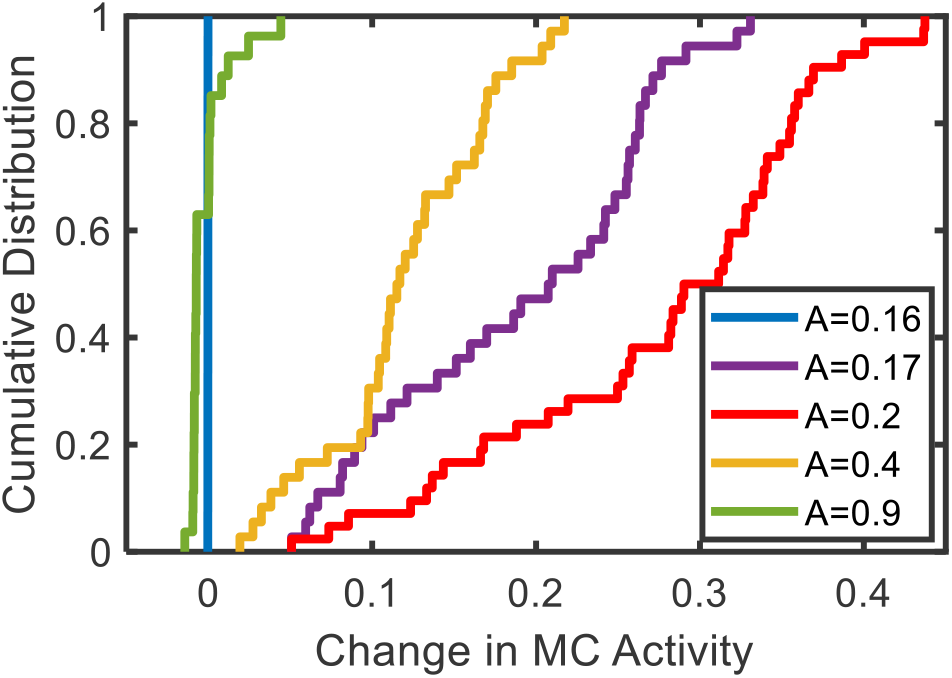
CDF for the increase in the MC response to odor *A* due to training with that odor at a reduced amplitude 2*A* after the network has been pre-trained with that odor at amplitude 2. For protocol see Fig.7.

### Forgetting due to Interference

**Fig S3.**
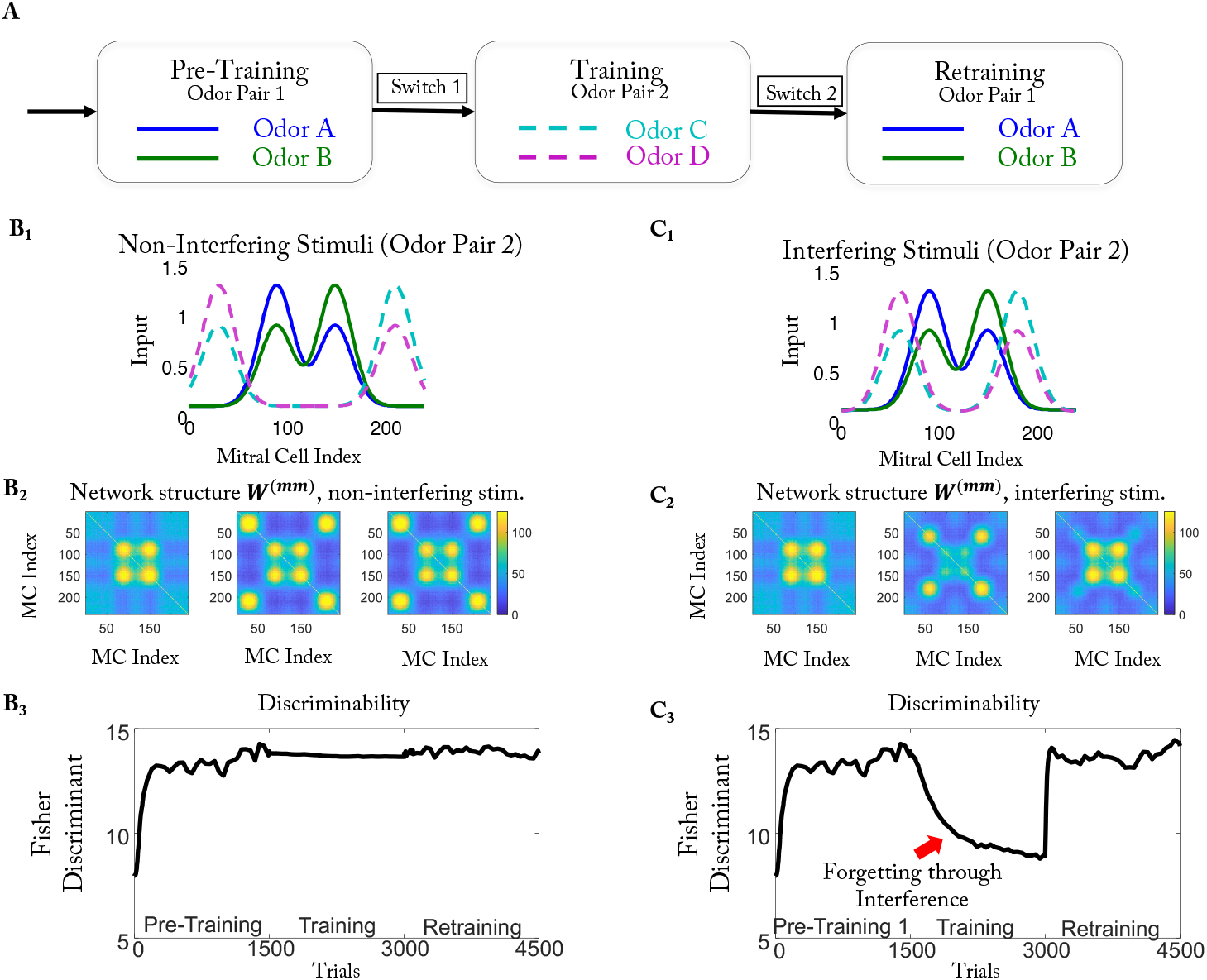
Interference leads to forgetting. (A) Training protocol. (B_1_) Non-interfering stimuli for odor pair 2. (B_2_) Connectivities after pre-training, training, and retraining, respectively. (B_3_) Discriminability is retained throughout the training. (C) as (B) for interfering stimuli. During the training the network forgets most of the previously learned structure. Retraining recovers the previous performance.

The behavior of the model depended strongly on the similarity of the two odor pairs (Fig S3B,C). When the MCs that were activated during the training did not overlap with those activated in the pre-training, the network preserved the previously learned structure (Fig S3B_2_). However, if there was significant overlap, the learning of the new stimuli interfered with the previously learned structure and that odor pair was forgotten during the training (Fig S3C_2_, middle panel). It was re-learned by re-training (Fig S3C_2_, right panel). The different evolution for non-interfering and for interfering stimuli is reflected in the Fisher discriminant ℱ^(1,2)^, which is given by the sum over the squares of the 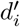 of the individual MCs in response to stimulus pair 1 and 2, respectively (see Methods). The Fisher discriminant ℱ^(1)^ remained high during the training with odor pair 2 if that pair was non-interfering (Fig S3B_3_), while it decreased if odor pair 2 was interfering (Fig S3C_3_). We expect that animals will spontaneously discriminate the odors in a pair only if their bulbar representations are sufficiently different, i.e., if ℱ is sufficiently large [38]. Thus, the model predicts that at the end of the training phase of a multi-phase perceptual learning task an animal will not spontaneously discriminate any more the odors it learned during pre-training if the training odors interfere with those used in the pre-training. However, for sufficiently different odor pairs, the spontaneous discrimination should remain intact.

### Rapid Relearning

Repeatedly switching between odor pairs did not impair the learning ability (Fig S4). In addition, the model predicts that re-learning during re-training is faster than the initial learning during pre-training and forgetting is slower than either of the learning processes.

**Fig S4.**
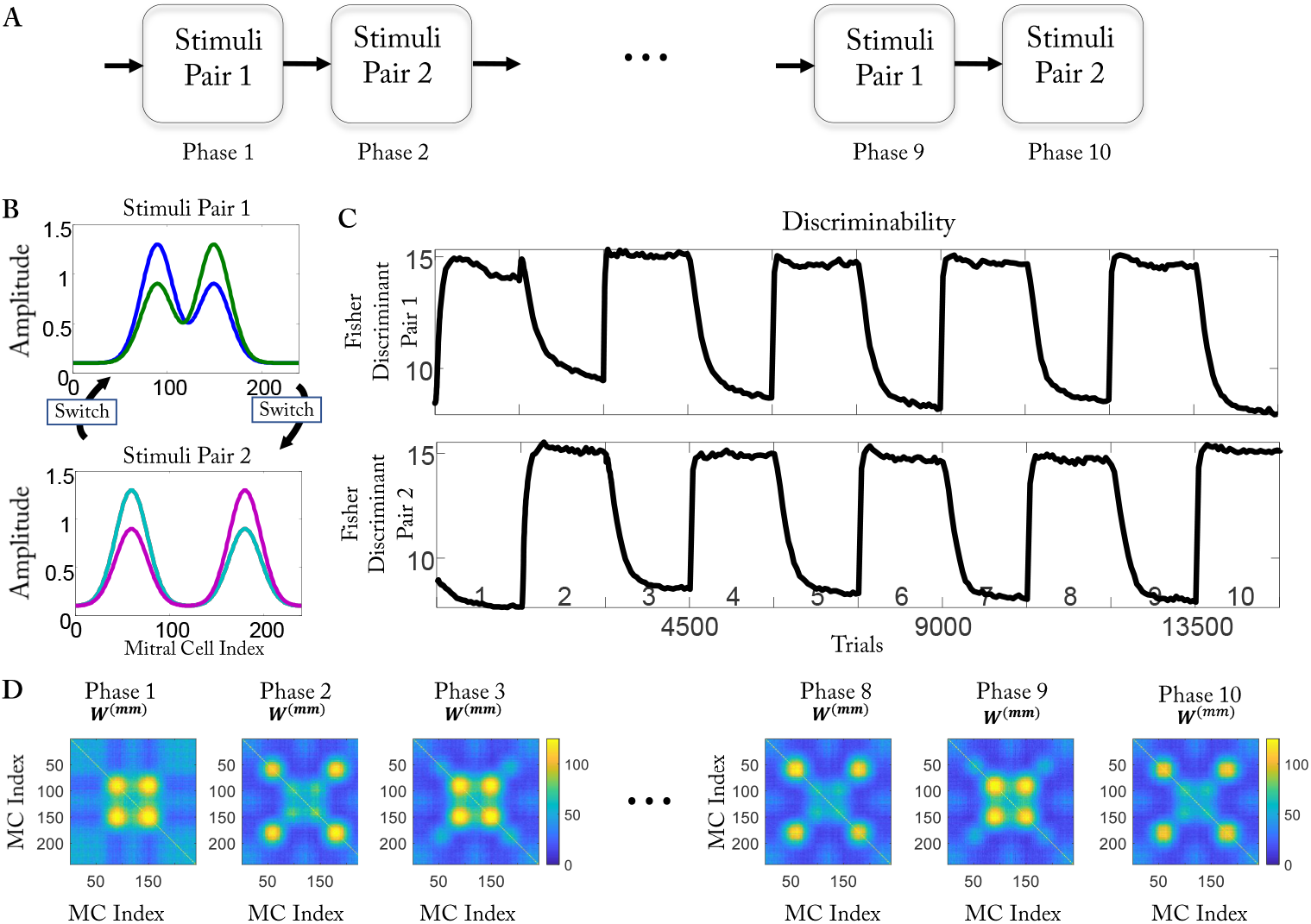
Alternating training does not impair learning ability. (A) Expanding the training protocol of S3A to 10 phases. (B) (Top) Stimuli of odor pair 1. (Bottom) Interfering stimuli of odor pair 2. (C) The Fisher discriminant of odor pair 1 (top) and pair 2 (bottom). Learning (increasing Fisher discriminant) proceeds faster than forgetting (decreasing Fisher discriminant). The numbers above the x-axis indicate the learning phase. (D) Effective connectivity *W* ^(*mm*)^ alternates.

### Resource Pool Model

**Fig S5.**
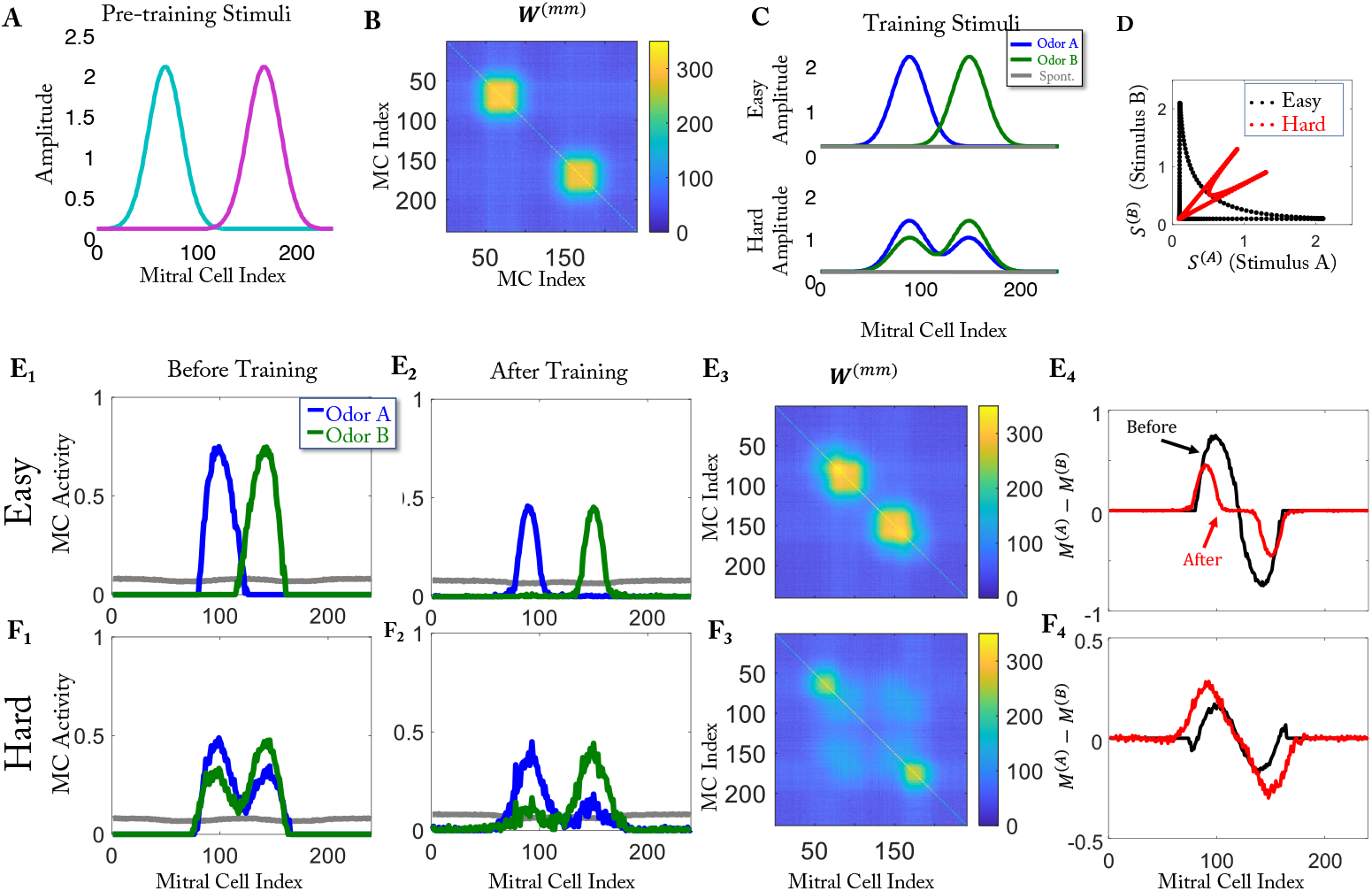
Results of the resource-pool model when trained with simplified stimuli. Here *γ* = 1.7 · 10^*-*3^. Training with dissimilar stimuli reduces their discriminability (E), whereas training with very similar enhances their discriminability (F) (cf. Fig.3)

### Impact of the Pre-Training

**Fig S6.**
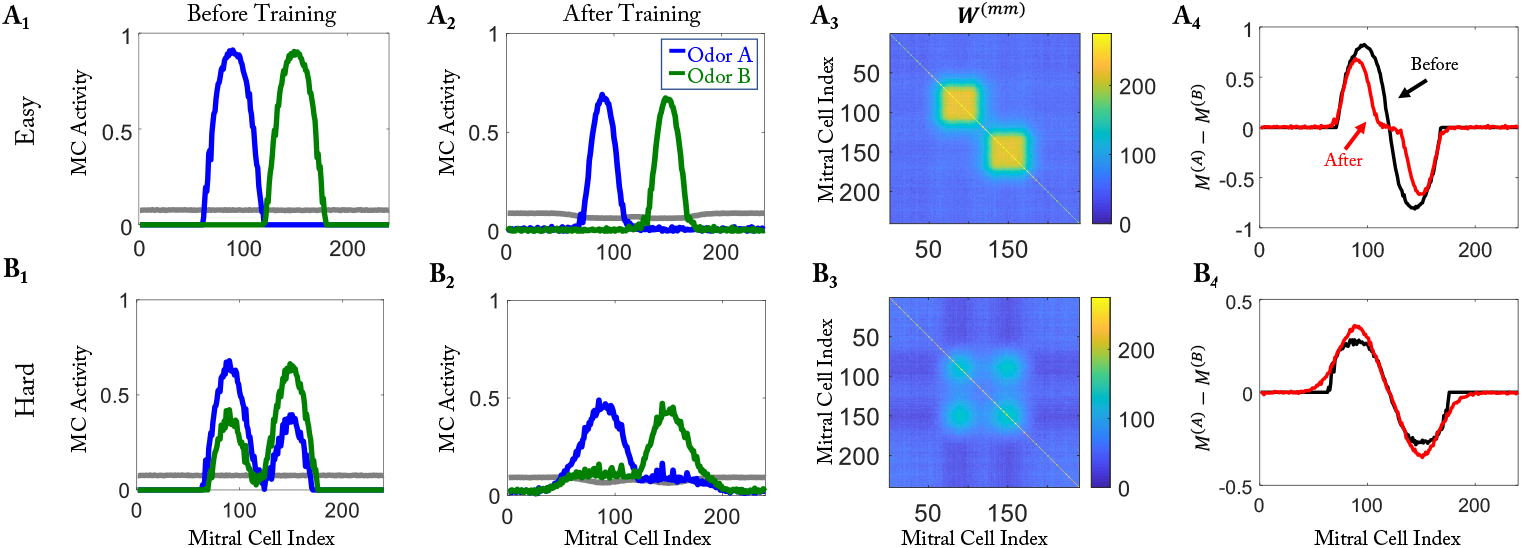
Training results without pre-training phase. The training starts with a random homogeneous network (cf. Fig 3)

### Robustness of the Model

To demonstrate the robustness of the model, we first assessed the impact of some modifications of the model.

For the formation of the reciprocal synapses little is known about the timescales of the maturation of the excitatory synapses relative to that of the inhibitory synapses. In the main part of the paper we assumed that inhibition becomes functional at the same as the excitation. Here, we tested if the model behaves differently if inhibitory synapses mature later than the excitatory synapses. If the inhibitory synapses become only functional 2 computational time steps after the excitatory synapses, the model performs similarly (Fig S7 A-C). Since 100 timesteps in our model correspond very roughly to about 1 day training in the experiments, a lag of 2 timesteps would correspond to a lag on the order of 30 minutes. However, the performance deteriorates if the lag is increased and the model fails for lags of 10 or larger. A detailed analysis of this break-down is beyond the scope of this study.

While action potentials can propagate along the secondary dendrites of MCs and can drive excitatory synapses on spines even far from the soma, the GC-driven inhibition originating from those spines is likely to affect the MC somata only within a quite limited spatial range. To assess the impact of this spatial limitation, we blocked the inhibition in half of the MC-GC pairs: while in all spines the MC→GC connections excited the GCs, only half of the MC←GC connections in those spines were effective in inhibiting the MCs. To make the total amount of inhibition comparable, we doubled the number of granule cells. With this modification, the results are similar to the original model (Fig S7 D, E).

**Fig S7.**
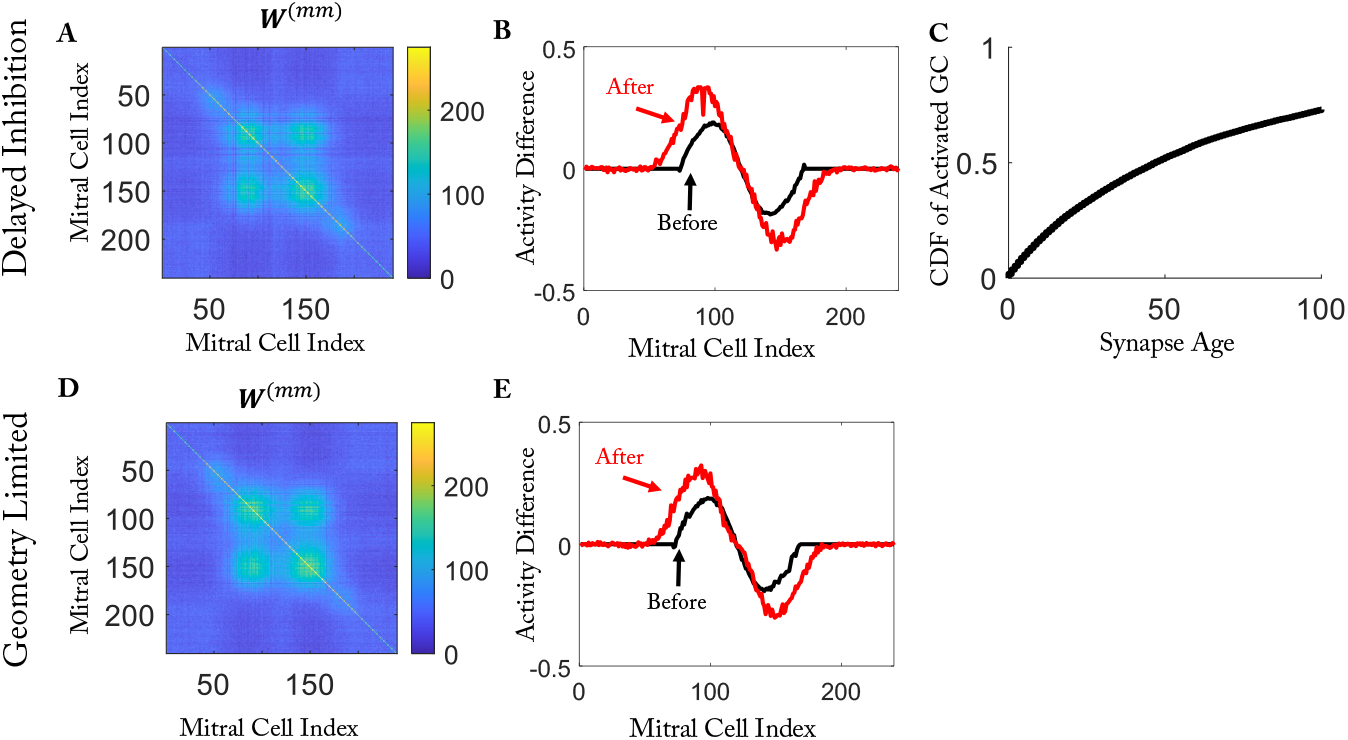
Modifications with asymmetric inhibitory and excitatory synapses show similar results in a hard task with simplified stimuli. (A - C) The results of setting maturation of the inhibitory synapses lagged 2 timesteps. (A) The effective connectivity after training. (B) Activity difference before and after training. (C) The cumulative distribution function of synapse age. When the lag is 2, only a limited fraction of connections does not inhibit MCs. (D, E) The results when the inhibitory synapses have a geometry limitation. Organized as (A, B).

Further, we tested the influence of various parameters on the model performance. Changing the total number *N*_*GC*_ of GCs affected the performance very little, since the number of GCs that were strongly activated and connected to specific MCs did not change significantly, which influenced the effective connectivity *W* ^(*mm*)^ only mildly (Fig S8). Analogously, changes in the inhibitory strength *γ* were largely compensated by a change in the number of activated GCs (Fig S9). This feedback stabilizing the overall inhibition arises from the reciprocal character of the MC-GC connections: new connections increase the inhibition of the MCs, which decreases the excitation of the GC and in turn inhibits the formation of new connections.

**Fig S8.**
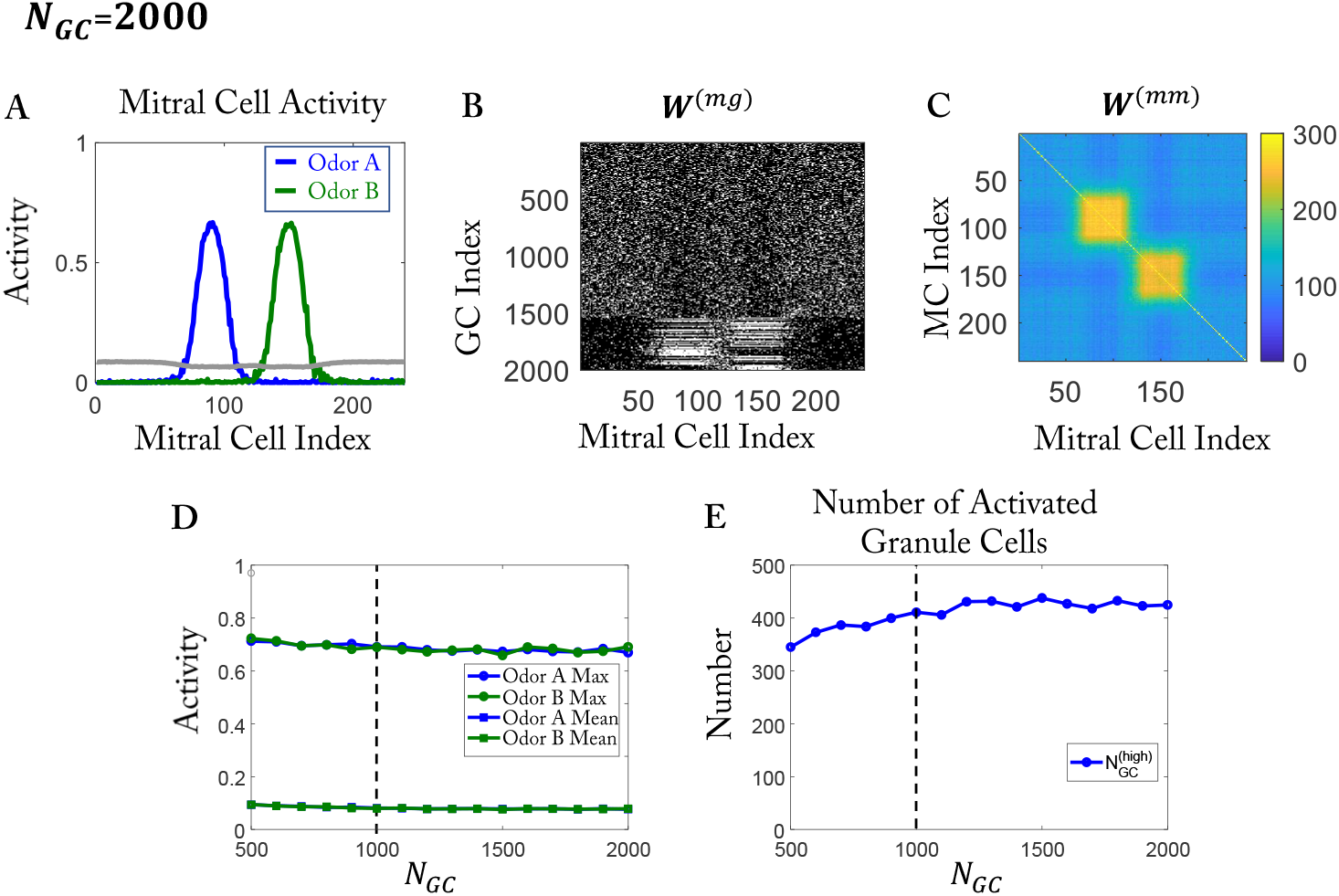
The overall inhibition of the model depends only weakly on the number *N*_*GC*_ of GCs. (A-C) Training with simplified easy stimuli as in Fig 1D and random initial network as in Fig 1F, but with twice as many GCs. (A) MC activity after training (cf. Fig 1H). (B) Connectivity *W* ^(*mg*)^ after training. (C) Effective connectivity *W* ^(*mm*)^. Compared to Fig 1J the connectivity is not quite as selective. (D) The maximal and mean MC activity after training as a function of the number of GCs. (E) The number of activated GCs (*G > G*^(1)^) depends only weakly on the total number of GCs. In (D, E) the vertical dashed line marks the value used in the rest of the paper.

What does control the overall inhibition, is the threshold *G*^(1)^ in the activation function *ϕ*(*G*) (Fig S10). Only GCs with activity larger than *G*^(1)^ form more synapses and generate stronger selective inhibition. Thus, for larger *G*^(1)^ fewer GCs had sufficiently large activity to sustain their synapses, which reduced the overall inhibition and increased the MC activity level. The change in overall inhibition could be associated with a change in the selectivity. To quantify this, we called a GC responsive to odor *A* (*B*), if after training its activity surpassed *G*^(1)^ for odor *A* (*B*). Despite the increase in the inhibition, the number of GCs that responded to both odors remained low when decreasing *G*^(1)^ (Fig S10H, red line), indicating that the selectivity depended very little on *G*^(1)^.

The other threshold in the activation function, *G*^(0)^, affects the removal but not the formation of synapses. For low values of *G*^(0)^ weakly activated GCs removed many of their synapses, particularly those connecting them to active MCs (Fig S11C, D). This lead to higher selectivity of the connectivity and with it to a weaker inhibition of the spontaneously active MCs that were not odor-driven (Fig S11B, E). It had, however, little impact on the maximal MC amplitudes (Fig S11H). The removal of synapses played a larger role in retaining and forgetting connections. To assess its impact, we trained the model in a second phase with a new pair of stimuli that overlapped with the stimuli learned in phase 1 (Fig S11I). For *G*^(0)^ = 0 any weak activity (0 *< G < G*^(1)^) triggered the removal of spines and lead in phase 2 to an almost complete forgetting of the connectivity learned in phase 1 (Fig S11D, K) and with it to poor discrimination of the previously learned odors (Fig S11N). However, for large *G*^(0)^ the previously learned connectivity was maintained (Fig S11G, M) and with it the discriminability of the previously learned stimuli. The discrimination of the newly learned stimulus pair 2 was, however, not significantly affected by *G*^(0)^ (Fig S11O).

**Fig S9.**
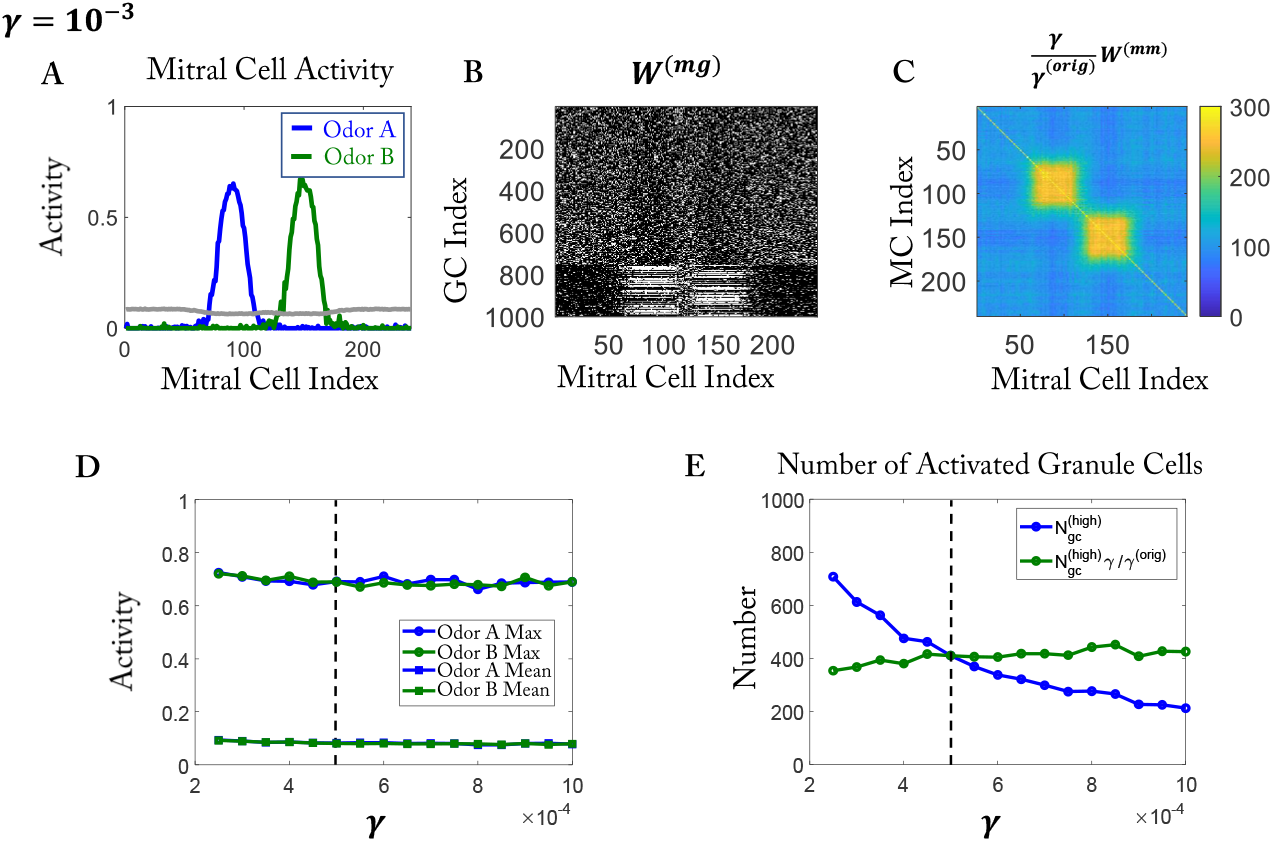
The overall inhibition of the model depends only weakly on the inhibitory strength *γ*. The results are organized as in Fig S8. (C) The connectivity is slightly less selective than in Fig 1J. (D,E) The vertical dashed line indicates the value used in the rest of the paper. (D, E) To allow a direct comparison, *W* ^(*mm*)^ and 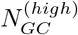 have been re-scaled by 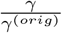 with *γ*^(*orig*)^ = 5 × 10^*-*4^.

The maximal number of connections *k* that each GC can make had a significant impact on MC amplitudes and on the selectivity of the connections. With increasing *k*, each GC integrated input from more MCs, and more easily surpassed the threshold *G*^(1)^. Each of these more activated GCs inhibited a larger number of MCs, increasing the overall inhibition (Fig S12C). When the maximal number of connections was larger than the number of MCs activated by one of the odors, not all connections of a GC could be made to MCs that were activated by odor *A*, say. The remaining connections could then be made to MCs corresponding to odor *B* without destabilizing the other synapses. As a result, GCs that responded to both odors emerged (Fig S12D).

**Fig S10.**
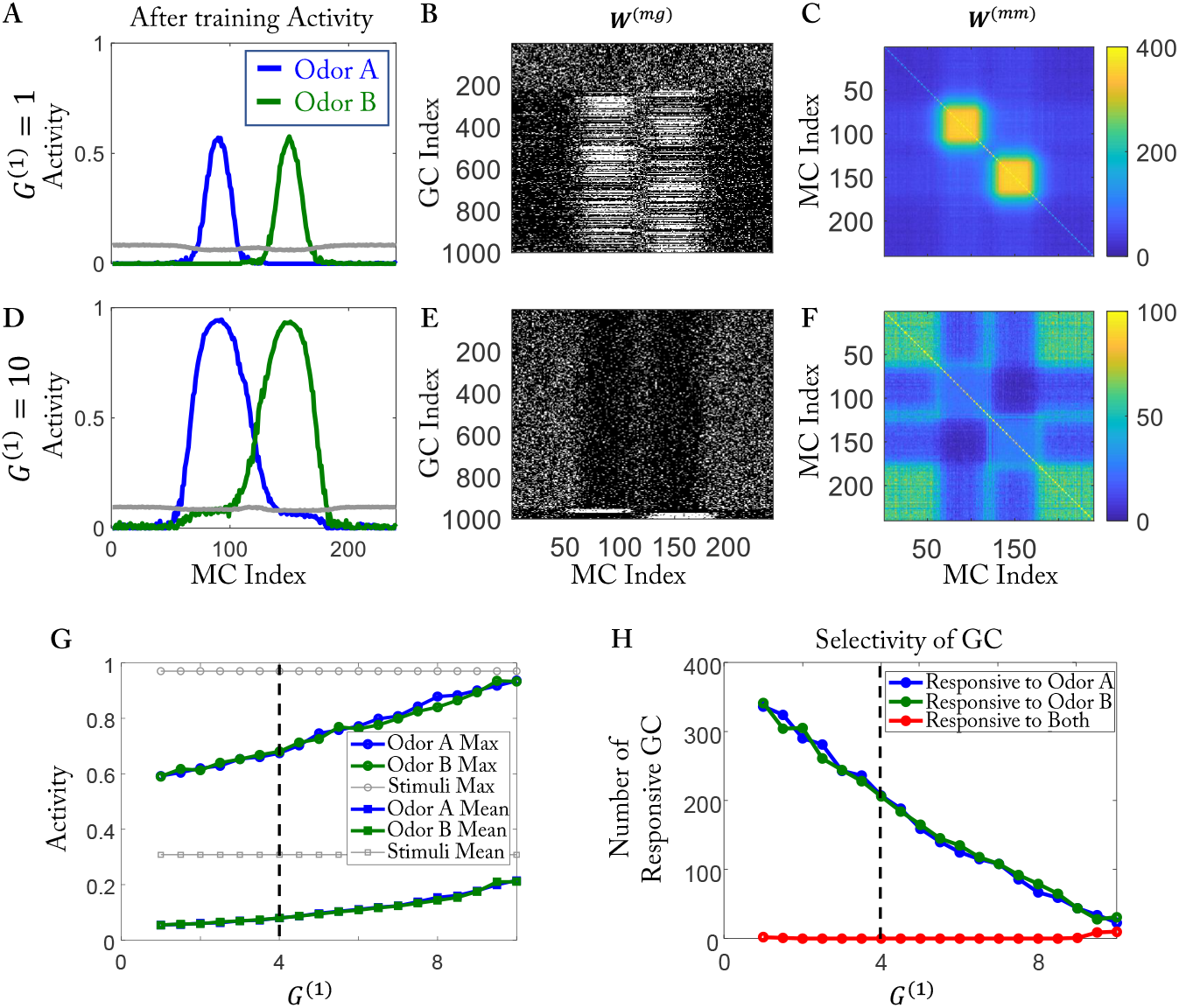
Increasing *G*^(1)^ impairs the ability to form connections between activated MCs and GCs. (A,B,C) Results for *G*^(1)^ = 1. (A) MC activity after training. (B) Connectivity *W* ^(*mg*)^ between MCs and GCs. (C) Effective connectivity *W* ^(*mm*)^. (D,E,G) as (A,B,C) except for *G*^(1)^ = 10. Activated MCs are connected with fewer GCs (compare E with B), resulting in weaker disynaptic inhibition (compare F with C) and higher MC activity (compare D to A). In (F) most of the GCs cannot reach the high threshold. As a result, the synapses that connect to strongly activated MCs are removed faster than those connecting to weakly activated MCs. Thus, the effective connectivity among the activated MCs is lower than the background. (G) Maximal and mean MC activity increase with increasing *G*^(1)^. The gray lines indicate the corresponding values without inhibition from GCs (by setting *γ* = 0). (H) The number of responsive GC decreases with increasing *G*^(1)^. (G,H) The vertical dashed line indicates the value used in the rest of the paper.

**Fig S11.**
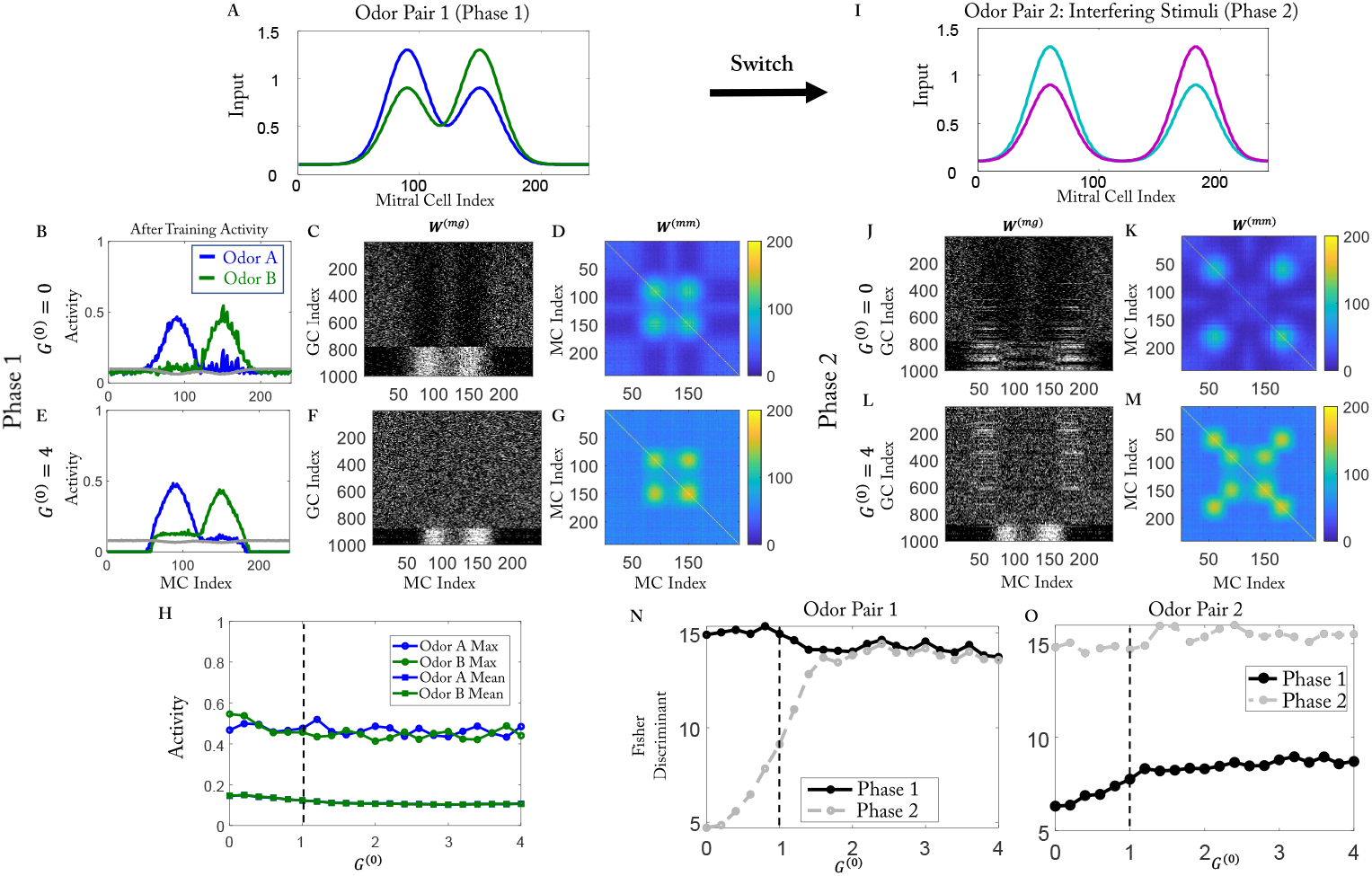
Learning ability is not affected by *G*^(0)^, but the memory is. (A-G) Phase 1. (A) Training stimuli in phase 1. (B) MC activity after training. (C) Connectivity *W* ^(*mg*)^. (D) Effective connectivity *W* ^(*mm*)^. (E to G) as (B to D) except for *G*^(0)^ = 4. (H) Maximal and mean MC activity after training in phase 1 as a function of *G*^(1)^. (I to M) as (A to G) but after Phase 2. (I) Interfering training stimuli. (J-M) For *G*^(0)^ = 0 the network forgets the previously learned connectivity, but not for *G*^(0)^ = 4. (N) The Fisher discriminant for odor pair 1 is unaffected by *G*^(0)^ in phase 1, but substantially reduced in phase 2 for small *G*^(0)^. (O) For odor pair 2 the Fisher discriminant depends only little on *G*^(0)^. The vertical dashed lines in (H,N,O) indicate the value used in the rest of the paper.

**Fig S12.**
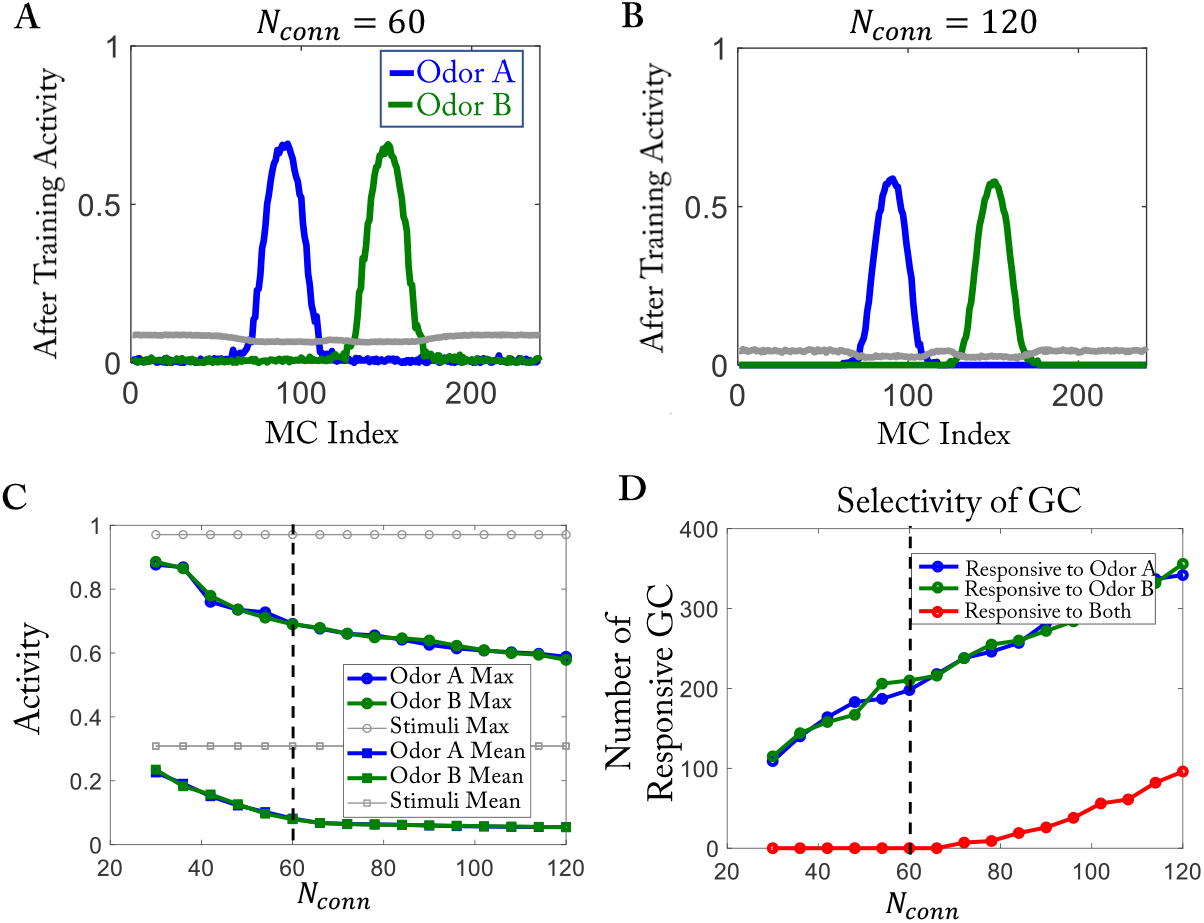
The selectivity of GCs is influenced by the number of connections each GC makes. Easy stimuli as in Fig 1D with random initial network as in Fig 1F. The ratio of maximal to initial number of connections is kept fixed at *k/N*_*conn*_ ≡ 1.1. (A, B) The MC activity is very similar for *N*_*conn*_ = 60 and *N*_*conn*_ = 120. (C) Maximal and mean MC activity decrease with increasing *N*_*conn*_. The gray lines indicate the corresponding values without inhibition from GCs (by setting *γ* = 0). (D) The selectivity of the GCs is impaired for *N*_*conn*_ *>* 60: a population of cells emerges that respond to both odors (red line).

**Fig S13.**
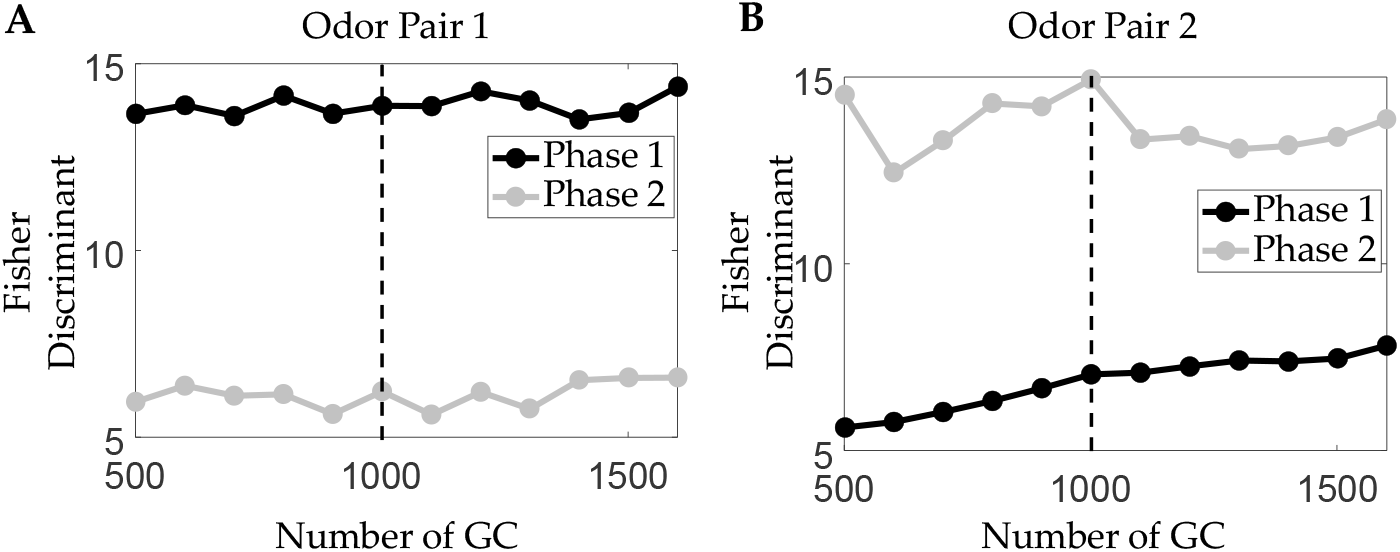
Increasing the number of GCs does not reduce the interference-induced forgetting of odor pair 1 during training phase 2. Independent of the number of GCs, the discriminability of odor pair 1, which is learned during phase 1 (black line in panel A), is substantially reduced by the subsequent training during phase 2 (gray line in panel A). The discriminability of odor pair 2 that is reached after training with those odors during phase 2 also does not depend on the number of GCs. Only the learning of odor pair 2 during the training with odor pair 1 is slightly improved in the larger network. (*G*^(0)^ = 1, cf. Fig S11N,O).

Dependence on the Threshold *θ*

**Fig S14.**
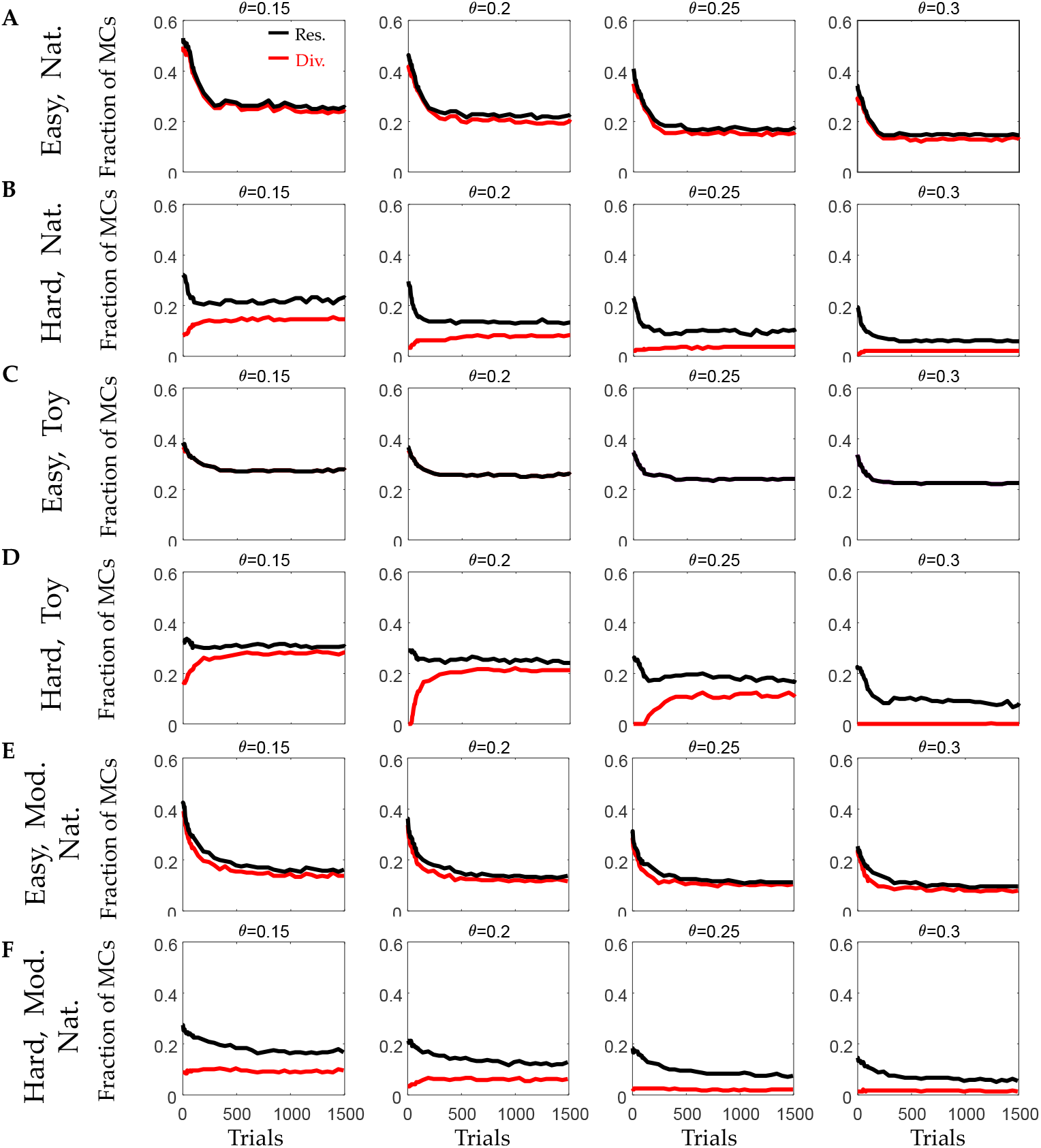
Qualitatively, the results for the fraction of responsive and divergent cells do not depend on the threshold *θ* for their classification. (A, B) Training with naturalistic stimuli for different thresholds *θ*. The results in Fig 2 and Fig 9 are based on *θ* = 0.2. (C, D) as (A, B) but for training with simplified stimuli. (E, F) as (A, B) but in the resource-pool model.

